# Changes in Higuchi’s fractal dimension across age in healthy human EEG are anti-correlated with changes in oscillatory power and 1/f slope

**DOI:** 10.1101/2024.06.15.599168

**Authors:** Srishty Aggarwal, Supratim Ray

**Author notes:** Declaration of interests: The authors declare no competing financial interests.

## Abstract

Non-linear dynamical methods such as Higuchi’s fractal dimension (HFD) are often used to study the complexities of brain activity. In human electroencephalogram (EEG), while power in the gamma band (30-70 Hz) and the slope of the power spectral density (PSD) have been shown to reduce with healthy aging, there are conflicting findings regarding how HFD and other measures of complexity vary with aging. Further, the dependence of HFD on features obtained from PSD (such as gamma power and slope) has not been thoroughly probed. To address these issues, we computed time and frequency resolved HFD for EEG data collected from elderly population (N=217), aged between 50-88 years, for baseline (BL) eyes open state and during a fixation task in which visual grating stimuli that induce strong gamma oscillations were presented. During BL, HFD increased with age for frequencies up to 150 Hz, but surprisingly showed an opposite trend at higher frequencies. Interestingly, this change in HFD was opposite to the age-related change in PSD 1/f slope. Further, stimulus-related changes in HFD were anti-correlated with the changes in oscillatory power. However, age classification using HFD was slightly better than classification using spectral features (power and slope). Further, stimulus and age-related changes in HFD persisted even after normalization with surrogates, showing the effect of non-linear dynamics on HFD. Therefore, HFD could be jointly sensitive to various spectral features as well as some non-linearities not captured using spectral analysis, which could enhance our understanding of brain dynamics underlying healthy aging.

## 1. Introduction

Aging is associated with a multitude of physical, biological, and cognitive transformations that exert significant impact on overall health and mental well-being. Throughout the transition from mid to late life, the brain undergoes notable age-related alterations in both morphology and physiological dynamics that are intricately intertwined with changes in cognition and behavior (Phillips and Andrés, 2010; Harada et al., 2013). In recent years, there has been a burgeoning interest in non-linear dynamics-based approaches to understand the complexity of brain activity (Anokhin et al., 1996; Drachman, 2006; Rabinovich et al., 2006; Lau et al., 2022; Díaz Beltrán et al., 2024). Fractal dimension (FD) has emerged as a powerful tool to probe non-linear changes in the brain and quantify the complexity and irregularity in neural processes by measuring fractal-like behavior in neurons and neural signals (Stam, 2005; Di Ieva, 2024). For example, its variation in neurons and inter-neuron connectivity is helpful in identifying different neurons and the range of their network cooperation (Smith et al., 2021).

There are a variety of methods used to compute FD, such as box counting dimension, correlation dimension (CD), Katz method and Higuchi’s method (Lau et al., 2022). Among these, Higuchi’s fractal dimension (HFD) (Higuchi, 1988; Esteller et al., 2001) has gained prominence as a complexity measure, leading to its recent recognition in different areas of neurophysiology (Werner, 2010) and for classification in neurodegenerative and neuromodulation studies including stress, depression, sleep, meditation and electrical stimulation (Kesić and Spasić, 2016; Kakumanu et al., 2018; Olejarczyk et al., 2024). Recently, it has been shown that HFD during resting state varies with age like an inverse parabola – it first increases from young to adulthood (up to 50 years) and then decreases (Zappasodi et al., 2015; Smits et al., 2016). These results suggest a reduction in complexity with age in elderly population, consistent with another study that used multiscale entropy (McIntosh et al., 2014), but inconsistent with other studies that have shown opposite trends in CD and entropy (Anokhin et al., 1996; Hogan et al., 2012). On the other hand, age related changes in electroencephalogram (EEG) measured using traditional linear methods like power spectral density (PSD) have shown more consistent trends. For example, oscillatory activity in alpha band (8-12 Hz) has been shown to reduce with aging (Babiloni et al., 2006; Scally et al., 2018). Similarly, slope of the PSD, which could indicate excitation-inhibition (E-I) balance, becomes shallower with age (Voytek et al., 2015). Stimulus-induced narrowband gamma oscillations, which are induced by presenting large visual gratings, also weaken with age (Murty et al., 2020).

Some studies have hinted at potential correlations between FD and oscillatory power (Kronholm et al., 2007; Akar et al., 2015; Stojadinović et al., 2020), but a detailed spectro-temporal comparison has not been done. As the literature on change of spectral measures with age is quite established, such associations between PSD measures and FD could help resolve the conflicting findings regarding the change in FD with age. Also, it could convey whether brain signals have significant non-linearity or are linear measures enough to provide detailed information about brain dynamics.

The primary objectives of this study are as follows:

i. **Examine the variation of HFD with respect to stimulus, aging and frequency:** We investigated the impact of visual grating stimuli on HFD and its modulation with age by utilizing EEG data that was previously collected from a large cohort of subjects (N=217) aged between 50-88 years while they viewed large gratings that induced gamma oscillations (Murty et al., 2020). Further, we explored the frequency dependence of HFD.
ii. **Assess the relationship between HFD and PSD measures:** As entropy and other non-linear measures have been shown to be strongly dependent on linear measures (Kosciessa et al., 2020; Päeske et al., 2023), we compared HFD variations with previously reported changes in stimulus-induced power and PSD slope with age (Murty et al., 2020; Aggarwal and Ray, 2023).
iii. **Investigate whether HFD provides additional information beyond PSD measures:** We used two approaches. First, we compared the performance of HFD versus PSD based measures for classification of the age of the subject (mid-aged versus old) from EEG data using linear discriminant analysis (LDA). Second, we performed surrogate analysis to test whether age and stimulus related changes in HFD are reflection of linear or non-linear changes in the signal.

## 2. Materials and Methods

### 2.1 HFD calculation

HFD was calculated using the algorithm proposed by Higuchi (1988). Here, for a time series *X*(1), *X*(2), *X*(3), …, *X*(*N*),, taken at a regular time interval, we first construct k new time series, 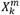, as

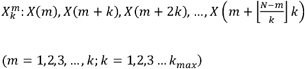

*m* and *k* are both integers and represent the initial time and the interval time respectively. The length *L*_*m*_(*k*) of the curve 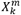, is given by

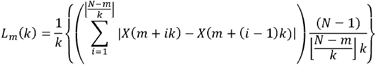

For each *k*, the length of the curve *L*(*k*) is then obtained by averaging the lengths *L*_*m*_(*k*), as

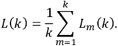

Finally, HFD is defined as

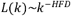

And obtained by computing L(k) over a range of k (from 1 to *k*_*max*_) and subsequently fitting a line through the log (L(k)) versus log (1/k) plot. We computed HFD using the Matlab code (https://in.mathworks.com/matlabcentral/fileexchange/124331-higfracdim).

HFD is best understood while considering a line (say with a slope of *m*). Average distance between two points separated by *k* is simply *mk*. Average “unit length” after dividing by *k* is simply *m*, and the total length (after accounting for the shortening of the time series) is therefore *m/k*. Hence, with increasing *k*, the total length should decrease as *k*^−1^, and hence a line has an HFD of 1.

HFD calculation is equivalent to a scenario where we embed the time series in boxes of various sizes (*1/k*) and compute the variation in the total length of the time series with the box size. A linear curve as HFD of one, but if the curve is more random, “zooming out” reduces the total length by a greater factor (and hence has a higher HFD). If the curve is really “dense” and covers the whole area on a planar graph, zooming out would lead to a reduction in length given by the square of the zooming factor, yielding an HFD of 2.

### 2.2 Synthetic Signals

To understand HFD and decide the optimum value of the parameter *k*_*max*_ for present analysis, we first computed HFD for synthetic signals. This included a straight line, a pure and a noisy sinusoid, and different forms of colored noise that yield fractal signals. A colored noise has power spectrum characterised by 1/*f*^*β*^, where *β* is the slope of the fractal signal. The slope is 0 for Gaussian noise or white noise. The slope between 1 and 3 defines a general class of temporal variability of fractals that is the fractional Brownian motion (fBm), with 2 as pure Brownian noise (Eke et al., 2000). fBm is a continuous-time Gaussian process *B*_*H*_(*t*) characterized by the “Hurst parameter” *H* (where 0 < *H* < 1), such that *B*_*H*_(*ct*) is statistically self-similar to *c*^*H*^*B*_*H*_(*t*) for a constant *c* > 0 (Mandelbrot and Van Ness, 1968). H determines the raggedness of the series, with a higher value leading to a smoother series. It is related to FD by FD = 2-H (Orey, 1970; Mandelbrot, 1985; Falconer, 2013). The autocorrelation function *R*(*τ*) of fBm is given by

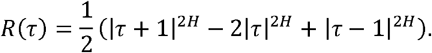

The PSD of fBm varies as *f*^−(2*H*+1)^ (Mandelbrot and Van Ness, 1968; Molz et al., 1997). Thus *β* = 2*H* + 1 = 5 – 2 *FD* (Higuchi, 1990)

While the increments in Brownian Motion (*X*(*t*) = *B*_*H*_(*s* + *t*) – *B*_*H*_(*s*)), for any *s*, are independent giving rise to Gaussian series, increments for fBm are not. If H < 0.5, then there is negative autocorrelation, i.e. if there is an increasing pattern in the previous steps, then it is likely that the current step will be decreasing whereas if H > 0.5, the autocorrelation is positive, indicating the opposite trend. Thus, H is an important index to predict the variation in time series and is widely used in stock market strategy as well (Matteo et al., 2005). We computed H using generalised Hurst exponent method (Barabási and Vicsek, 1991; Matteo et al., 2005; Mandelbrot, 2013), as described in the Supplementary Data section.

### 2.3 Neural Signals

The detailed explanation of experimental setup and data collection have been provided in the earlier studies (Murty et al., 2020, 2021; Kumar et al., 2022; Aggarwal and Ray, 2023), which we summarize below.

#### 2.3.1 Dataset

We analyzed the EEG dataset from the Tata Longitudinal Study of Aging, comprising 236 cognitively healthy individuals (132 males, 104 females), aged 50-88 years (Murty et al., 2020). Participants were recruited from metropolitan areas of Bengaluru, Karnataka, India. A team of psychiatrists, psychologists, and neurologists at the National Institute of Mental Health and Neurosciences (NIMHANS) and M.S. Ramaiah Hospital in Bengaluru clinically assessed and diagnosed them as cognitively healthy. The assessment included standard cognitive tests such as Addenbrooke’s Cognitive Examination-III (ACE-III), Clinical Dementia Rating (CDR) scale, and Hindi Mental State Examination (HMSE). To ensure data quality, 10 participants were excluded due to noise (see Artifact Rejection subsection). For 9 participants, data was recorded using 32 channels instead of the standard 64-channel setup, which made it difficult to average across subjects for some analyses (such as topoplots). These subjects were also removed, as done in our previous studies on this dataset (Aggarwal and Ray, 2023; Kumar and Ray, 2023). This resulted in a final dataset of 217 participants (119 males, 98 females) with usable 64-channel EEG recordings. Analogous to the previous studies (Murty et al., 2020, 2021; Kumar et al., 2022; Aggarwal and Ray, 2023), participants were divided into two age categories: 50–64 years (mid; N=90) and >64 years (old; N=127).

All participants gave informed consent before taking part in the study and received monetary compensation. The study procedures received approval from the Institutional Human Ethics Committees of Indian Institute of Science, NIMHANS, and M.S. Ramaiah Hospital, Bengaluru.

#### 2.3.2 Experimental Settings and Behavioural task

EEG data were collected using 64-channel active electrodes (actiCap) with the BrainAmp DC EEG acquisition system (Brain Products GmbH). The electrode placement followed the international 10–10 system and was referenced online to FCz. The raw signals underwent online filtering, with a passband between 0.016 Hz (first-order filter) and 1 kHz (fifth-order Butterworth filter), and were sampled at 2.5 kHz with a 16-bit resolution (0.1 μV/bit). Participants were instructed to sit in a dark room facing a gamma-corrected LCD monitor (BenQ XL2411; dimensions: 20.92 × 11.77 inches; resolution: 1280 × 720 pixels; refresh rate: 100 Hz), with their heads stabilized using a chin rest. The monitor was placed at a distance of 58 ± 0.7 cm (mean ± SD; range: 54.9–61.0 cm) from the participants. Eye movements were monitored using an Eyelink 1000 system (SR Research Ltd), with a sampling rate of 500 Hz.

The task was a passive visual fixation task involving the full screen grating stimuli. It contained a single session that spanned around 20 minutes and was divided in two-three blocks with 3–5 minutes breaks in between, based on participants’ comfort. A fixation spot (0.1°) was shown at the centre of the screen during the trial, on which the subjects were asked to fixate. After an initial 100 ms blank period, two to three full screen grating stimuli along with the fixation dot were shown for 800 ms with an interstimulus interval of 700 ms. The stimulus presentation was controlled using a customized software running on MAC OS. Full contrast sinusoidal luminance achromatic gratings with one of three spatial frequencies (1, 2, and 4 cycles per degree) and 4 orientations (0°, 45°, 90°, and 135°), chosen pseudorandomly, were presented as the stimuli.

#### 2.3.3 Artifact Rejection

We implemented a fully automated artifact rejection procedure as outlined in (Murty and Ray, 2022; Aggarwal and Ray, 2023). Eye blinks or any changes in eye position outside a 5° fixation window from −0.5 s to 0.75 s following stimulus onset were excluded offline, resulting in the rejection of 16.4 ± 14.2% (mean ± SD; Mid:14.4 ± 13.0%, Old: 17.9 ± 14.8%, p = 0.08, tstat = −1.78, t-test) of repeats. Electrodes with impedance greater than 25 kΩ were discarded, leaving the final set of electrodes with an impedance of 5.48 ± 1.83 kΩ (Mid: 5.57 ± 1.92 kΩ, Old: 5.47 ± 1.75 kΩ, p = 0.69, tstat = 0.40, t-test). For the remaining electrodes, outliers were identified as repeats with deviations from the mean signal by more than 6 SD in either the time or frequency domains, and electrodes with more than 30% outliers were excluded. Additionally, repeats from visual electrodes (P3, P1, P2, PO3, POz, PO4, O1, Oz, O2) or more than 10% of other electrodes that were deemed bad were also discarded, resulting in a set of common bad repeats for each subject. In total, 15.58 ± 5.48% (Mid: 15.43 ± 5.63%, Old: 15.63 ± 5.49, p=0.79, tstat = −0.26, t-test) repeats were rejected. Next, we calculated the power spectrum slopes between 56 Hz and 84 Hz for each unipolar electrode, rejecting electrodes with negative slopes, showing an unconventional upward tilt in the power spectrum.

After removing bad electrodes and repeats based on the above criteria, we calculated the root mean square (RMS) value of the time series for all remaining trials for each electrode. Repeats with RMS values outside the range of 1.25 µV to 35 µV were considered new outliers. Electrodes with more than 30% new outliers were excluded, and any new outlier found in visual electrodes or common to more than 10% of other electrodes was classified as a bad repeat and added to the list of existing bad repeats. Following these steps of rejection of bad electrodes (common across all blocks) and repeats, 55.20 ± 6.68 (Mid: 56.17 ± 6.62 Old: 54.52 ± 6.66, p = 0.07, tstat = −1.80, t-test) electrodes and 295.53 ± 67.40 (Mid: 305.40 ± 62.74, Old: 288.54 ± 69.92, p =0.07, tstat = 1.82, t-test) repeats per subject remained for the final analysis. The artifacts were not different significantly between the two age groups.

We also discarded blocks without at least one good unipolar electrode from each of the following electrode groups: left visual anterolateral (P3, P1, PO3, O1), right visual anterolateral (P2, P4, PO4, O2), and posteromedial (POz, Oz). The remaining good blocks for each subject were pooled for final analysis. This led to the exclusion of 10 subjects who did not have any analyzable blocks.

#### 2.3.4 EEG Data Analysis

All data analyses were conducted using custom codes written in MATLAB (MathWorks. Inc; RRID:SCR_001622). The analyses were done using unipolar reference scheme (i.e., all signals referenced to a single electrode, FCz, which was how the data was originally recorded). We used −0.5 to 0 s before and 0.25 to 0.75 s period after the stimulus onset as the “baseline (BL)” and “stimulus (ST)” periods, respectively. To study the effect of stimulus and aging, we filtered the signals between 1 and 90 Hz. For frequency dependent analysis, the signals were filtered in different frequency ranges as described in Sections 3.1.2.3 and 3.2. The filtered signals were obtained using an infinite filter response fourth order Chebyshev type I band pass filter with bandpass ripple of 0.2 Hz. They were also filtered for the line noise at 50 Hz and its harmonics within ± 2 Hz using the infinite filter response fourth order Chebyshev type I band stop filter.

##### 2.3.4.1 HFD analysis

For EEG data, HFD was obtained for each trial for BL and ST time series with 1250 data points and then averaged across trials for each electrode. The time-frequency HFD plots were generated using the non-overlapping 0.1 s time window between −0.25 and 1.15 s and 12 Hz frequency window between 1 to 109 Hz. Narrower windows (<12 Hz) resulted in diminished visibility of HFD-related effects in the TF plots due to smoothening of the signals and suppression of non-linear characteristics by narrowband filtering.

HFD is dependent on *k*_*max*_. Values of *k*_*max*_ between 6-10 have been shown appropriate for HFD computation (Accardo et al., 1997; Spasić et al., 2005). We chose *k*_*max*_ = 10 for low frequency signals (<100 Hz), since the log (L) vs log (1/k) curve was linear over this range. However, at higher frequencies, the log (L) vs log (1/k) curve became non-linear for *k*_*max*_ = 10. So, to maintain uniformity, we chose *k*_*max*_ = 5 while analysing the frequency dependence of HFD.

##### 2.3.4.2 Power and Slope Analysis

We obtained power spectrum and time-frequency power spectrogram using the multi-taper method with a single taper using the Chronux Toolbox ((Bokil et al., 2010), RRID:SCR_005547) for individual trials and then averaged across the trials for each electrode. The spectrograms were obtained using a moving window of size 0.25 s and a step size of 0.025 s, yielding a frequency resolution of 4 Hz. The change in power (in dB) is calculated as

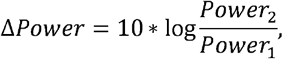

where Power_1_ and Power_2_ is power of the group 1 (“BL” and “Old” in case of stimulus and aging comparisons respectively) and group 2 (“ST”/ “Mid”) respectively.

We calculated the slope of the 1/f aperiodic component of the trial-averaged PSD for each electrode using the Matlab wrapper for Fitting Oscillations and One Over f (FOOOF) toolbox (Donoghue et al., 2020). Similar to (Aggarwal and Ray, 2023), we set the parameters for FOOOF model as: peak width limits: [4 8]; minimum peak height: 0.2; maximum number of peaks: 5; peak threshold: 2.0; and aperiodic mode: ‘fixed’. Slopes were computed in frequency ranges from 4 to 350 Hz in steps of 20 Hz with centre frequency between 40 Hz and 300 Hz, and the frequency width 100 Hz. To avoid poor fitting, we deemed ± 4 Hz around the peaks associated with the line noise at 50 Hz and its harmonics as “noise peaks” and excluded these ranges if these noise peaks lay near the end points of frequency range to fit. Further, we excluded slopes less than 0.01 from further analysis.

The correlation coefficients (*ρ*) and p-value for the change in HFD vs the change in power and slope were obtained using the Matlab command *corrcoef*. We used *fitlm* function in Matlab that generates the model parameters: *β*1, *β*2 corresponding to *y* = *β*1 + *β*2 * *x*, to obtain the regression line.

##### 2.3.4.3 Power matching

To remove the dependence of parameter changes (here band power and slope) on HFD, we followed the method as described in (Churchland et al., 2010; Kumar and Ray, 2023). We sub-selected subjects from each group such that change in parameter had the same distribution within the groups. For this, we first computed the distribution of subjects with respect to the parameter in each group. Then, we matched the number of subjects from each group within each parameter bin by randomly selecting subjects from the group with more subjects, to match the subjects in the other group. Ultimately, the subjects in each bin were adjusted to arrive at the common distribution for the two age groups. As each iteration produced a different subset of subjects, the above procedure was repeated 10 times to obtain average value of HFD for the parameter matched groups.

##### 2.3.4.4 Classifier analysis

We predicted the age group (mid or old) using a classifier based on linear discriminant analysis (LDA). Both spectral and HFD measures were considered as distinct features. We assessed the classifier performance on the features through the area under the receiver operating characteristic (ROC) curve (AUC) on 5-fold cross-validation sets, where the ROC curve represents the true positive rate against false positive rate. We used the Matlab command *fitdiscr* to generate the LDA based classifier model and then *perfcurve* to compute AUC for the features.

##### 2.3.4.5 Surrogate Analysis

The main purpose of the surrogate analysis was to generate synthetic data which had the same PSD and amplitude distribution as the original data. We obtained the surrogates of the filtered data for each trial using two methods: phase randomization (Theiler et al., 1992) and iterated amplitude adjusted Fourier transform (IAAFT) (Schreiber and Schmitz, 1996). We implemented these methods using Chaotic Systems Toolbox https://in.mathworks.com/matlabcentral/fileexchange/1597-chaotic-systems-toolbox.

In phase randomization, surrogates were generated by keeping the spectral coefficients of the original time series but randomizing the phases. It preserved the power spectrum but distorted the amplitude distribution in time domain, which can produce spurious identification of non-random structure (Rapp et al., 1994).

On the other hand, IAAFT has been found to be more effective to determine the presence of non-linearity in signals. Briefly, it begins by generating the phase randomized surrogates. As step (i), a new surrogate is obtained by rank ordering the original time series with respect to the old surrogate, which better aligns the phases between the two time series. In the next step (ii), the new surrogates are generated by applying inverse Fourier transform to the product of the power coefficients of the original time series with the phase of the surrogates generated in step (i). Steps (i) and (ii) are then iterated until the error difference between the power spectrum of the original time series and surrogates reduces. The generated surrogate by IAAFT closely matches the original time series in terms of both the amplitude distribution and the power spectrum.

#### 2.3.5 Electrode Grouping

Scalp maps were created using the topoplot function of EEGLAB toolbox ((Delorme and Makeig, 2004), RRID:SCR_007292) with standard *Acticap 64* unipolar montage. Following (Aggarwal and Ray, 2023), all the electrodes were divided into 5 groups as occipital (O; O1, Oz, O2, PO3, PO4, PO7, PO8, PO9, PO10, POz), centro-parietal (CP; CP1, CP2, CP3, CP4, CP5, CP6, CPz, P1, P2, P3, P4, P5, P6, P7, P8, Pz), fronto-central (FC; FC1, FC2, FC3, FC4, FC5, FC6, C1, C2, C3, C4, C5, C6, Cz), frontal (F; Fp1, Fp2, F1, F2, F3, F4, F5, F6, F7, F8, Fz, AF3, AF4, AF7, AF8) and temporal (T; T7, T8, TP7, TP8, TP9, TP10, FT7, FT8, FT9, FT10). FCz was used as the reference electrode and Fpz served as the ground.. To observe the stimulus effect, a subgroup of occipital and centro-parietal electrodes (P3, P1, P2, PO3, POz, PO4, O1, Oz and O2) were chosen, in which strong stimulus-induced gamma was observed and termed as high priority electrodes (HP) (Murty et al., 2020).

### 2.4 Statistical Analysis

Wilcoxon signed rank test was used to compare HFD and H between BL and ST across all trials and all subjects. Kruskal-Wallis (K-W) test and Wilcoxon rank sum (WRS) test were used to compare the medians for power, slope and HFD across groups, as done in previous studies (Murty et al., 2020; Aggarwal and Ray, 2023). The standard error of median (SEM) was computed after bootstrapping over 10,000 iterations. To account for multiple comparisons across electrode groups, we employed false discovery rate (FDR) correction using the Benjamini-Hochberg procedure (Benjamini and Hochberg, 1995).

## Data and Code Availability

All spectral analyses were performed using Chronux toolbox (version 2.10), available at http://chronux.org. Slopes were obtained using matlab wrapper for FOOOF (https://github.com/fooof-tools/fooof_mat). The codes for HFD were taken from https://in.mathworks.com/matlabcentral/fileexchange/124331-higfracdim. Surrogate analysis was done using Chaotic Systems Toolbox (https://in.mathworks.com/matlabcentral/fileexchange/1597-chaotic-systems-toolbox). The violin plots are plotted using the ‘gramm data visualization toolbox’ (Morel, 2018). The codes and HFD data for generating figures in the present work are available at https://github.com/4srihy/TLSAEEGProjectPrograms/tree/master/HFD_AgeProjectCodes. Raw data, downsampled to 250 Hz, is freely available at https://osf.io/ebryn/.

## 3. Results

### 3.1 Variation of HFD in synthetic and neural signals

#### 3.1.1 Synthetic Signals

To illustrate HFD, we first generated synthetic signals with 1000 data points with known properties and computed their HFD (with *k*_*max*_ =10, see Methods for details), as depicted in Fig. 1A. To compare the slope of log (L) vs log (1/k) curve, we changed the offset for each condition such that all traces had the same value of log (L) at k = 1 (or log (1/k) = 0; right panel of Fig. 1A). Each condition was simulated 10 times and mean and SD were computed.

**Fig. 1:**
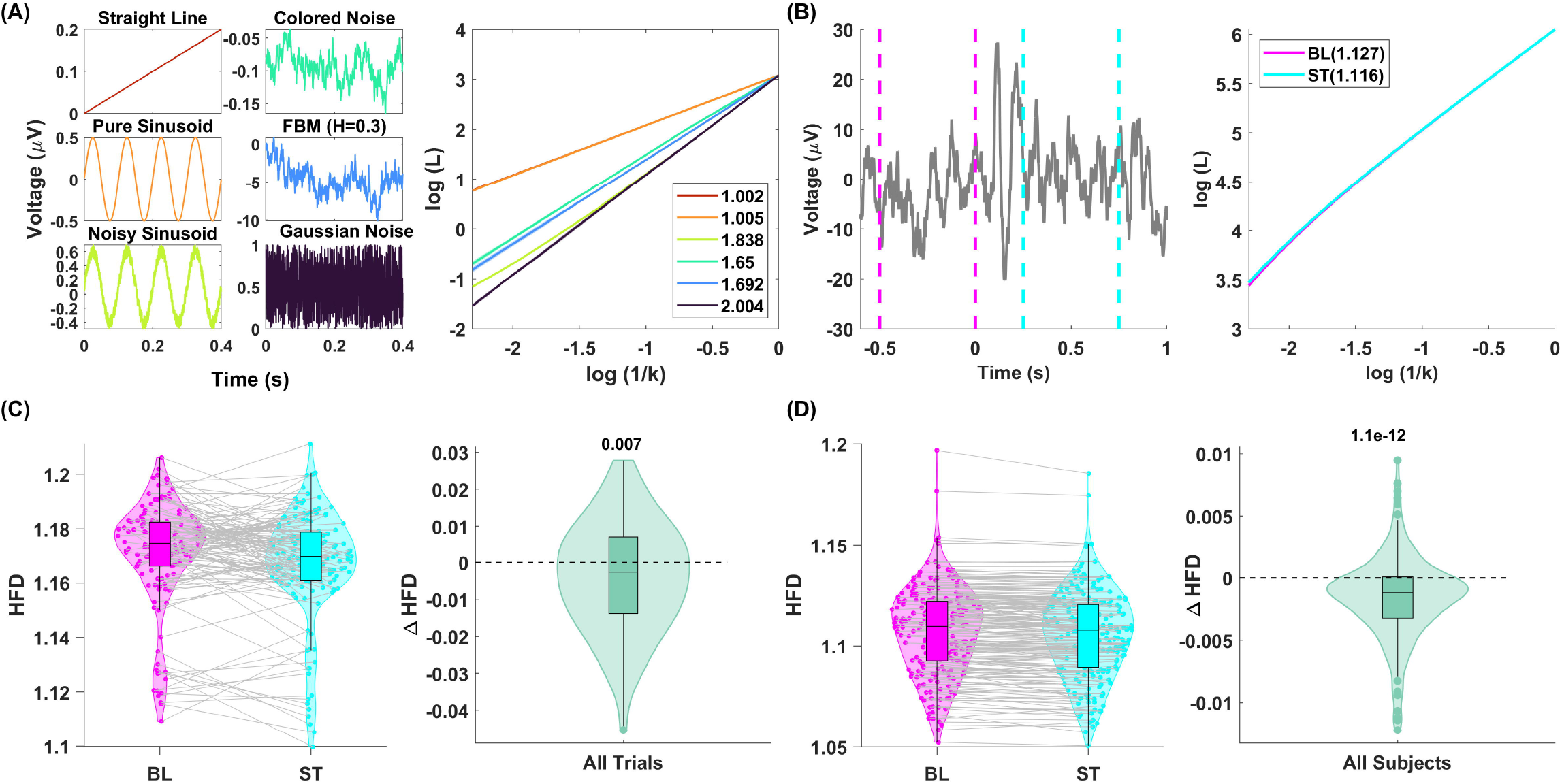
**(A)** Different synthetic signals (left panel: straight line of slope 0.5 (brown); sine wave of frequency 10 Hz (orange); sine wave with 40% Gaussian noise added (light green);colored noise with slope (*β*)of 1.7 (dark green); fractional Brownian motion (fBm) with Hurst index (H) of 0.3 (blue) and Gaussian noise (violet)) and corresponding average log (L) vs log (1/k) curves (right panel). The shaded region around the curves shows ± SD, computed by running the simulations 10 times (not visible for most traces since these SDs were very small). The legend shows the average HFD for these signals. **(B)** Time series for one trial for one subject for one occipital electrode (Oz), filtered between 1 to 90 Hz (left panel). Magenta and cyan vertical lines indicate the BL and ST periods, respectively. Log (L) vs log (1/k) curves for the filtered baseline (BL) and stimulus (ST) segments with *k* _*max*_ = 10 (right panel). The HFD (slope of the curves) values are indicated in the legend. **(C)** Violin plots with matched pairs showing the distribution of HFD for BL and ST (left panel) and ΔHFD (right panel) for all trials (n = 139) for the electrode Oz of the one sample subject. The p-value for median computed from Wilcoxon signed rank test is indicated on the top in the right panel. **(D)** Violin plots showing the distribution of HFD for BL and ST (left panel) and Δ HFD (right panel) for all subjects (N=217) for the high priority (HP) electrode group.

For a straight line (brown trace), the HFD was (mean ± SD) 1.00 ± 0.00, as expected (see Methods for details). Even for a pure sinusoid (frequency of 10 Hz; orange trace), HFD remained close to 1 (1.00 ± 0.00), with the log (L) vs log (1/k) plot overlapping with the straight-line condition. Since HFD depends on the roughness in the curves, it is ~1 for all straight lines and low frequency pure sinusoids. Adding 40% Gaussian noise to the sinusoid (light green trace) increased HFD to 1.84 ± 0.01. Though the overall shape of the noisy sinusoid remained the same as the pure sinusoid, the added fluctuations which can be seen as increased thickness of the light green trace as compared to the orange trace led to a higher HFD. For colored noise with *β* of 1.7 (dark green trace), HFD and H were 1.65 ± 0.03 and 0.34 ± 0.02, consistent with the theoretical values of FD and H of 1.65 and 0.35 (see Methods for details). For fBm with H parameter set to 0.3 (blue trace), HFD was 1.69 ± 0.02, consistent with theoretical FD of 1.7. For Gaussian noise covering nearly the full graph (violet trace), HFD was 2.00 ± 0.01.

#### 3.1.2 Neural signals (EEG)

Next, we examined EEG data for 217 healthy adults aged between 50 and 88 years, grouped as mid (50-64 years; N=90) and old (>64 years; N=127). Subjects performed a fixation task in which full-screen gratings were presented for 800 ms with an inter-stimulus interval of 700 ms. We considered −500 to 0 ms before and 250 to 750 ms periods after stimulus onset as baseline (BL) and stimulus (ST) periods respectively and computed HFD. We filtered the EEG signals in different frequency ranges using an infinite filter response fourth order Chebyshev type I band pass filter with bandpass ripple of 0.2 Hz. Further, to remove the line noise at 50 Hz and its harmonics, we filtered the signals at corresponding frequencies within ± 2 Hz using the infinite filter response fourth order Chebyshev type I band stop filter.

##### 3.1.2.1 Stimulus presentation reduced HFD

We first probed the effect of stimulus on HFD. Figure 1B illustrates the process of computation of HFD for a single trial for one electrode (Oz) for one subject. We took the raw time series and filtered it between 1-90 Hz (Fig. 1B, left panel). We computed the HFD for the BL (−0.5 to 0 s) and ST (0.25 to 0.75 s) segments, as highlighted in Fig. 1B left panel, as slopes of log (L) vs log (1/k) curves, shown in Fig. 1B right panel. The curve for BL was slightly steeper than for ST, leading to lower HFD for ST than BL, though the difference was quite small. However, this difference was consistent across trials for this electrode and subject, as shown in Fig. 1C left panel. The median HFD difference between ST and BL for all trials (n = 139) for a single block was significantly lower than 0 (p = 0.007, zval = 2.708, signed rank = 6153, Wilcoxon signed rank test), as depicted in Fig. 1C, right panel. The visual stimulus mainly changed the activity in the occipital and other nearby electrodes. Therefore, we chose a subgroup of these electrodes, termed “high-priority” (HP) electrodes here (highlighted in Fig. 2A) and in previous studies (Murty et al., 2020, 2021) to study the effect of stimulus in detail. Across the total population of 217 subjects, although the HFD values varied considerably, the reduction between BL and ST averaged across all trials for HP electrode group was consistent, such that the difference was highly significant (Fig. 1D; p =1.1 × 10^−12^, zval = 7.120, signed rank = 18419, Wilcoxon signed rank test).

**Fig. 2:**
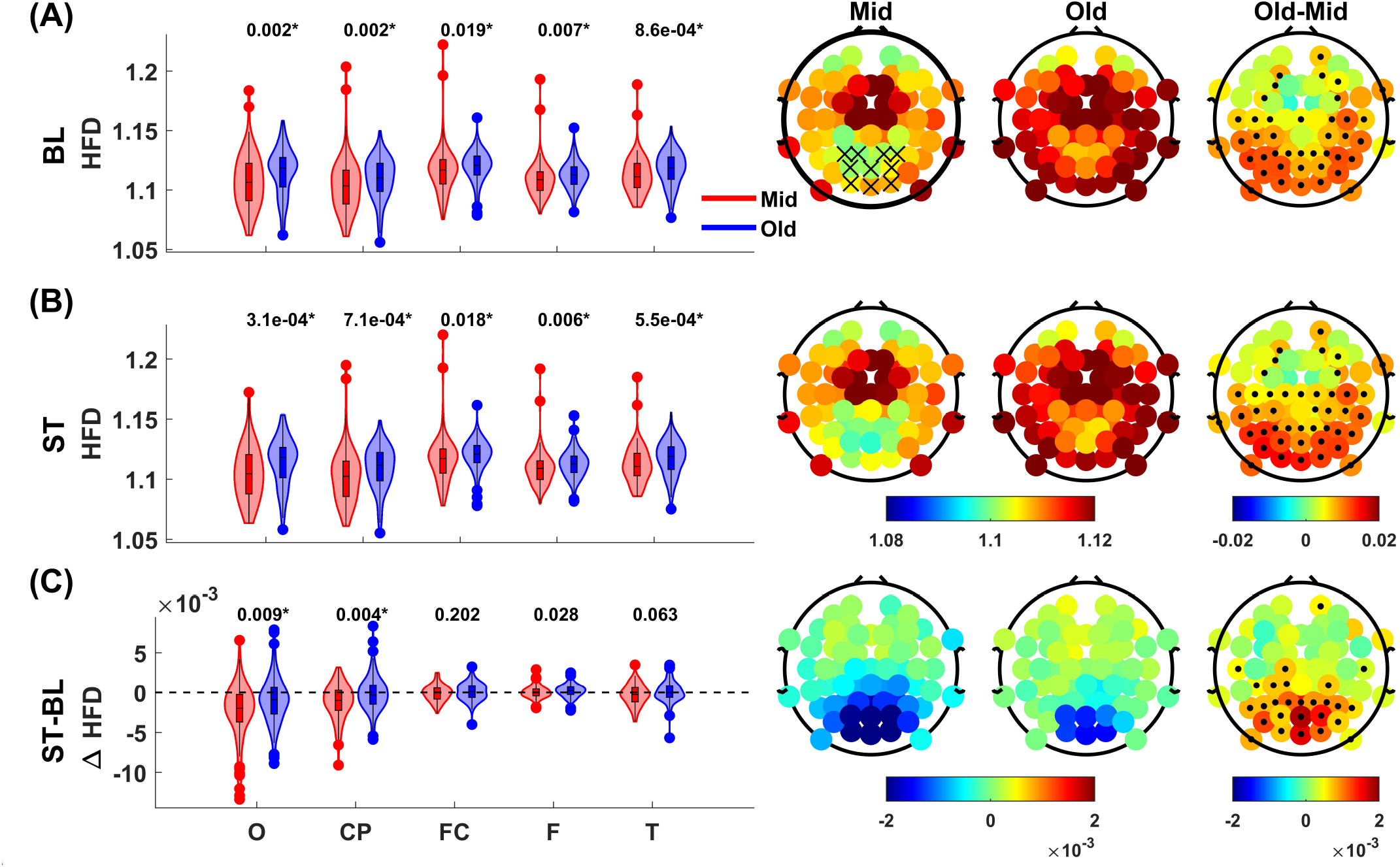
**(A)** (Left) Violin plots showing the comparison of HFD for BL in mid and old aged group for different electrode groups. The p-values, computed from Wilcoxon rank sum (WRS) test are indicated at the top of the distribution for the corresponding electrode group. * indicates that the p-value remain significant after false discovery rate (FDR) correction. (Right) Topoplots showing the median HFD across the electrodes for mid (left), old (center) and difference: old-mid (right). ‘x’ in the left topoplot indicate high-priority (HP) electrodes. Electrodes with significant difference (p < 0.05, WRS test) are highlighted in black in the right topoplot. Similar comparison is shown for ST in **(B)** and Diff (ST-BL) in **(C)**.

To cross-verify the HFD computation and the choice of *k*_*max*_, we also checked the effect of stimulus on the Hurst Index (H; Supplementary Fig. 1), which has been associated with long-range correlations (Hurst, 1951; Shang, 2020). Though H is theoretically related to FD as FD = 2-H (Orey, 1970; Mandelbrot, 1985; Falconer, 2013), for real signals, the relation may not be satisfied (Amezquita-Sanchez et al., 2019). We found that H increased with stimulus, thus, showing an opposite trend of HFD. Moreover, the sum of HFD and H tended to 2 (1.984 ± 0.001 for both BL and ST), which to our knowledge has not been observed for real EEG signals. This essentially validates the choice of *k*_*max*_ and use of HFD as the intrinsic fractal dimension of the signals. Since the relation between H and HFD was maintained for all our signals, we have limited our further analysis to the changes in HFD only.

##### 3.1.2.2 HFD increased with age

Next, we divided EEG electrodes into 5 groups (see Methods for details) and computed HFD for BL and ST periods. Subjects were divided into mid (50-64 years) and old (≥ 65 years) age groups. We computed HFD for the filtered signal between 1 to 90 Hz with *k*_*max*_ set to 10. For BL, HFD was significantly higher for old group than mid in all electrode groups (p-values shown in the plot, WRS test, FDR corrected) as shown in Fig. 2A violin plot and topoplots. The statistics for violin plot comparisons in Fig. 2A-C are summarized in Supplementary tables 1A-C. The HFD difference between the two age groups was significant for a wide range of *k*_*max*_(5-35) suggesting that these results did not depend on a particular choice of *k*_*max*_.

While HFD was higher in central electrodes than occipital electrodes for both age groups (Fig. 2A left and centre topoplots), the difference in HFD was higher in occipital electrodes and gradually decreased towards the frontal region, as depicted in Fig. 2A right topoplot (old-mid). Similar difference was observed for ST period (Fig. 2B). However, the change in HFD between ST and BL was more negative in mid group as compared to the old group mainly in occipital (p = 0.009, zval = −2.630, ranksum = 8611, WRS test, FDR corrected) and centro-parietal regions (p = 0.004, zval = −2.876, ranksum = 8499, WRS test, FDR corrected; Fig. 2C).

##### 3.1.2.3 HFD increased with frequency

As shown in previous studies, the visual stimuli induced strong stimulus-induced gamma oscillations that weakened with age (Murty et al., 2020). To test whether changes in HFD were related to the changes in the power spectrum of the signals, we computed HFD as a function of frequency of the signal. Figure 3A shows the BL segment of the signal (the full signal is shown in the inset) for a single trial for Oz electrode for one subject. We then filtered the signal in different frequency ranges as indicated in Fig. 3B using an infinite filter response fourth order Chebyshev type I band pass filter with bandpass ripple of 0.2 Hz. We chose these ranges to avoid the line noise and its harmonics. The corresponding log (L) vs log (1/k) curves up to *k*_*max*_ =10 with dotted lines are shown in Fig. 3C. As the frequency range increased, the log (L) curve became more non-linear. To maintain uniformity for choosing linear range for log (L) vs log (1/k), we computed HFD for all frequency ranges up to *k*_*max*_= 5, as indicated by solid lines in Fig. 3C. Figure 3D shows the variation of the HFD with centre frequency for the frequency ranges shown in Fig. 3B. As expected, HFD increased with frequency since higher frequency range signals were less smooth. We further calculated the HFD for shorter segments of the signal to generate the time-frequency (TF) HFD profile. Figure 3E depicts this profile for all trials (n = 139) for HP electrode group for one subject. The corresponding delta TF plot in Fig. 3F, obtained after subtracting the mean BL HFD values at each frequency, showed a strong reduction in HFD from 1 to 40 Hz during the transient period (0 to 0.25 s) and a steady decrease around ~20 Hz during the ST period. As shown later, this reduction in HFD after stimulus onset matches with the increase in oscillatory power in the slow gamma range after stimulus onset.

**Fig. 3:**
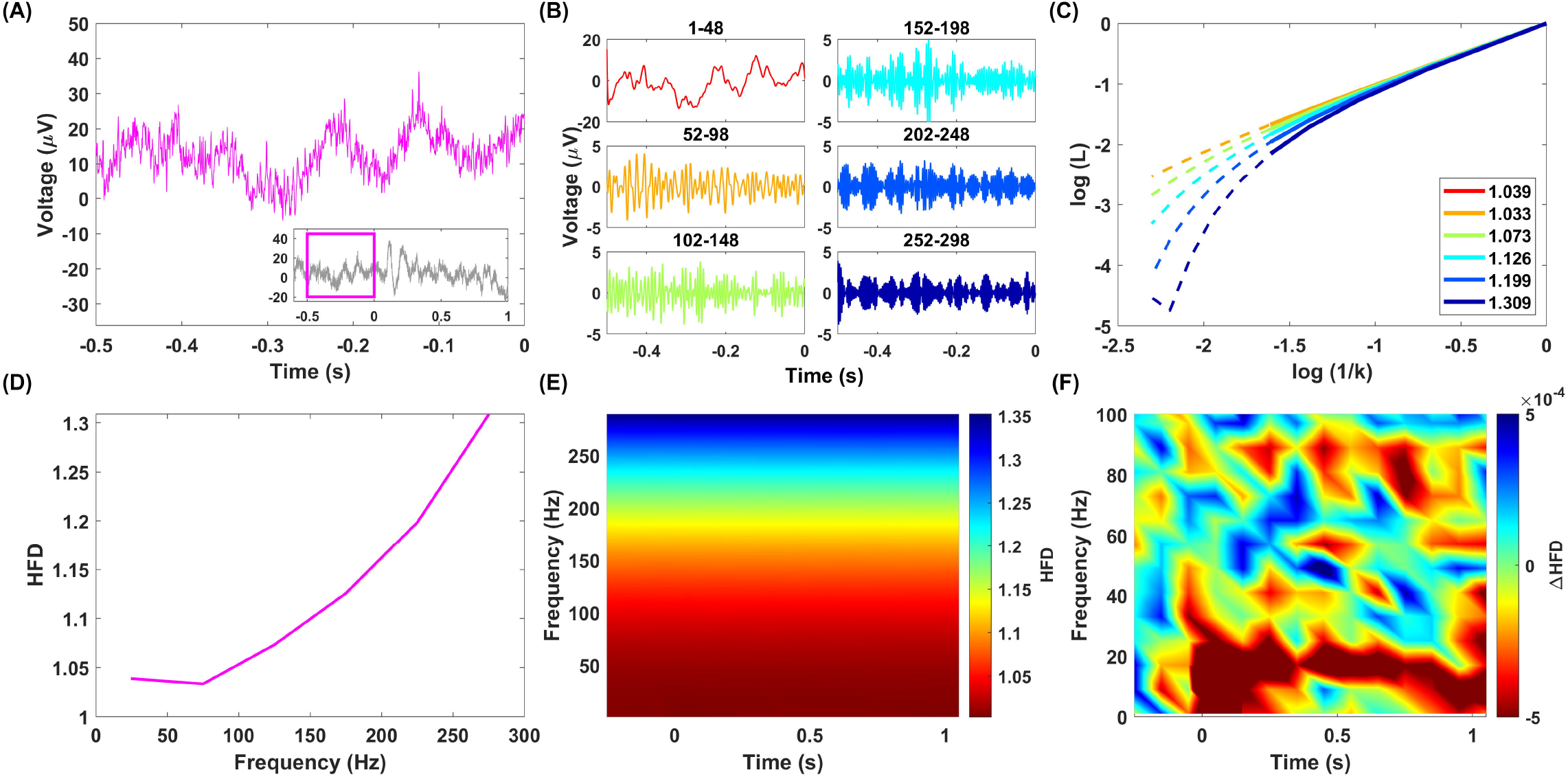
**(A)** BL time series for a single trial for one electrode (Oz) from one subject, segmented from the whole time series as shown in the inset with the box highlighting the BL period. **(B)** BL time series filtered in different frequency ranges. **(C)** Corresponding log (L) vs log (1/k) curves for *kmax* =10(dotted) and *k*_*max*_ =5(solid). The HFD computed with *k* _max_=5are indicated in legends in (C) and are plotted with respect to centre frequency in **(D). (E)** Time-frequency (TF) plot for HFD averaged across all trials (n = 139) for HP electrode group. **(F)** Corresponding delta TF plot showing the change in HFD during stimulus and transient periods with respect to baseline.

### 3.2 Dependence of HFD on Spectral Features

#### 3.2.1 Change in frequency dependent HFD with age during baseline was opposite to the change in the slope of the PSD

Next, we explored the changes in HFD with age as a function of frequency. We computed HFD (with *k*_*max*_ = 6 for signal in the BL period after filtering with centre frequencies from 40 Hz to 300 Hz (in steps of 10 Hz) by taking ± 50 Hz around each centre frequency (for centre frequencies of 40 and 50 Hz, the lower limit was set to 4 Hz). Figure 4A left panel shows the HFD as a function of frequency in HP electrode group for mid and old age groups. Although the difference of HFD between these groups was small, the curves were significantly different at many frequency ranges, as indicated by black (p<0.05; K-W test) and green (p<0.01; K-W test) bars. The change in HFD (old-mid) is shown in Fig. 4A right panel. Interestingly, the age-dependent increase in HFD during BL, as shown in Fig. 2A was present only up to ~150 Hz. At higher frequencies, HFD was lower for old group as compared to mid group.

**Fig. 4:**
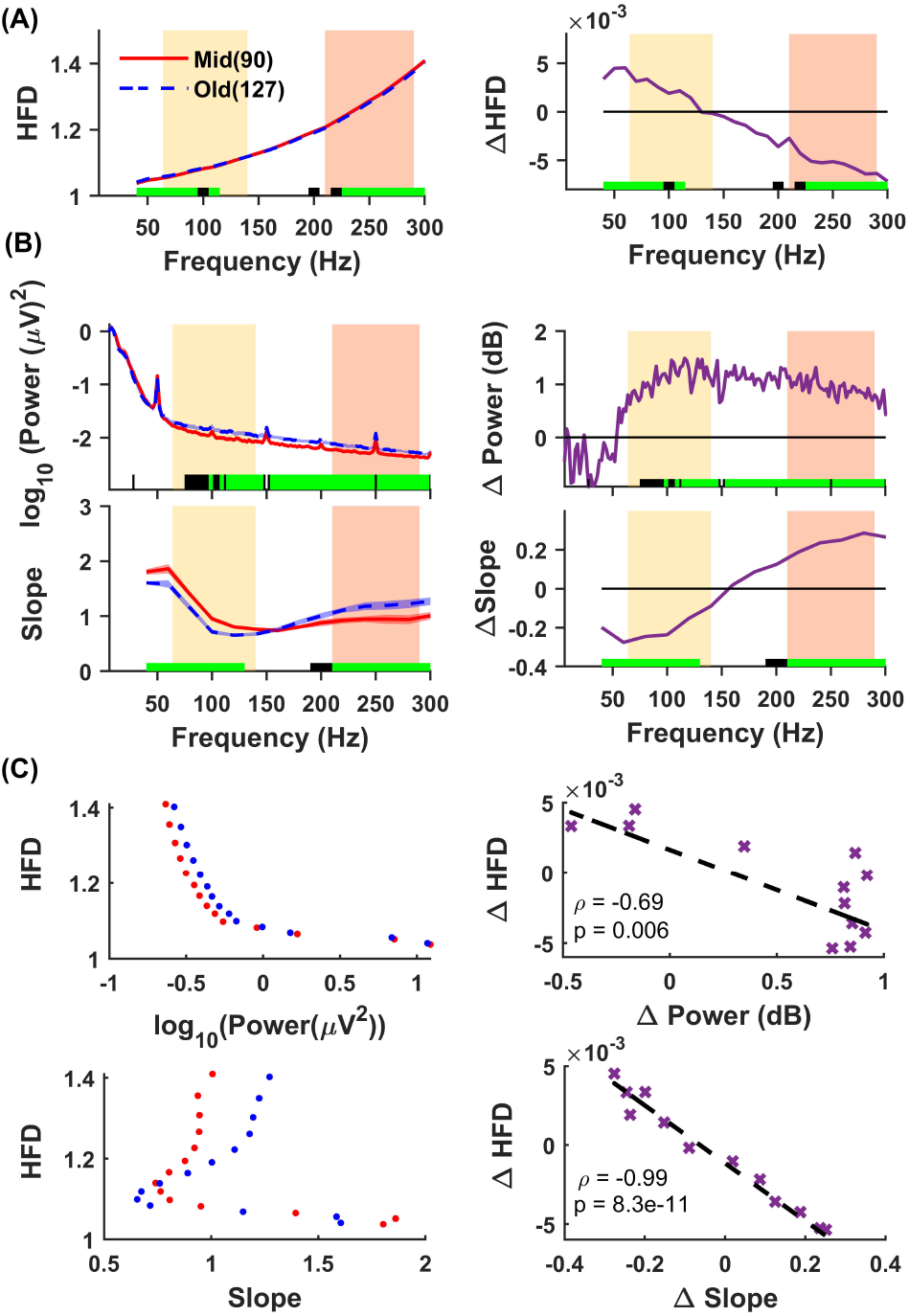
**(A)** HFD, **(B) (Top)** Power spectral density (PSD), **(bottom)** Slope versus frequency for BL period in HP electrodes for mid (solid) and old (dash) groups (left) and their corresponding difference (old-mid; right). In the left panels, solid traces represent the median and shaded region around them indicates ± Standard Error of Median (SEM) across subjects, computed after bootstrapping over 10,000 iterations (shaded regions are barely visible due to low SEMs). Coloured bars at the abscissa at the bottom represent significance of differences between mid and old (black: p < 0.05 and green: p < 0.01, Kruskal-Wallis (K-W) test). Yellow and orange boxes represent the low frequency range (LFR; 64-140 Hz) and the high frequency range (HFR; 210-290 Hz) respectively. In the right panels, violet traces represent the difference of median of old and mid group shown in the left panels and coloured bars at the abscissa are same as the left ones. **(C) (Left)** HFD versus power (top) and slope (bottom) at the centre frequencies between 40-300 Hz in steps of 20 Hz and frequency width of 100 Hz (left) for mid and old. **(Right)** The corresponding group difference of Δ HFD vs ΔPower (top) and Δ Slope (bottom) respectively. ‘x’ represents the data points and the regression line is in black. The correlation coefficient () and p-value are indicated in each subplot.

To probe the dependence of HFD on spectral measures, we compared the age-related BL HFD changes with the similar changes in power and slope. We plotted the BL PSD for mid and old groups (Fig. 4B top left panel) with frequency resolution 2 Hz and the change in power in dB (Fig. 4B top right panel). We also plotted the slope and the corresponding change with age (old-mid) computed between the frequency ranges from 4 to 350 Hz in steps of 20 Hz with centre frequency between 40 Hz and 300 Hz and the frequency width of 100 Hz. While old group had lower power at frequencies till ~50 Hz and higher power beyond (Fig. 4B top panel), slope was lower for old-aged group up to ~150 Hz and then was higher (Fig. 4B bottom panel), as reported previously (Aggarwal and Ray, 2023). The trend of the HFD change seemed opposite to the age dependent slope change but not much on the change in power.

To better quantify these changes, we plotted HFD with power and slope for the two age groups and corresponding ΔHFD with ΔPower and ΔSlope. Here power is computed by summing the PSD for frequency ranges obtained at the centre frequencies between 40 and 300 Hz in the steps of 20 Hz with the frequency width of 100 Hz. Figure 4C top panel shows the dependence of HFD on power for mid and old (left) and the change in power (in dB) versus the change in HFD between the groups (right). As HFD increased with frequency due to the increase in roughness of signal and power decreased with frequency, HFD and power were anti-correlated. There was a significant negative correlation between ΔHFD and ΔPower (correlation coefficient (*ρ*)= −0.69, p = 0.006). On the other hand, while HFD did not show a specific relationship with slope as depicted in Fig. 4C bottom left panel, ΔHFD showed a surprisingly strong negative correlation with ΔSlope with age (Fig. 4C bottom right panel; *ρ* = −0.99,= 8.3 × 10^−11^). This anti-correlation between ΔHFD and ΔSlope persisted even if *k*_*max*_ was not kept the same for all frequency ranges (data not shown). Thus, the change in the HFD in aging as a function of frequency strongly reflected the corresponding changes in PSD slope, indicating that HFD depends more on the rate of change of power with frequency than the magnitude of change of power itself.

#### 3.2.2 Change in frequency dependent HFD with age during stimulus was opposite to the change in Power

To probe age-related effect of ST on HFD and its dependence on gamma power, we first compared the ΔPower between the two age groups with respect to frequency (Fig. 5A top panel). While there was no significant power difference in the alpha range (8-12 Hz; pink band), mid-aged group had significantly higher power than old group in both slow (purple, 20-34 Hz) and fast (grey, 36-66 Hz; p < 0.05: black and <0.01: green, K-W test). The same was visible in TF spectra for change in power obtained by subtracting average BL power from the raw power in Fig. 4B top panel for mid (left), old (centre) and old-mid (right), as shown previously (Murty et al., 2020). To compare corresponding changes in HFD, we computed ΔHFD (ST-BL) for continuous non-overlapping frequency ranges with centre frequencies lying between 7 and 103 Hz and frequency width of 12 Hz (Figure 5A, bottom plot), and for non-overlapping 0.1 s time window between −0.25 and 1.15 s and frequency width of 12 Hz for TF HFD plots (Fig 5B bottom panels). While ΔHFD with respect to stimulus was higher in alpha range, it showed a reduction in gamma bands. In the alpha band, ΔHFD was not different between the two age groups. However, ΔHFD was significantly more negative in mid-aged group than old in slow and fast gamma bands (colorbars of the TF plot for ΔHFD are reversed compared to ΔPower to observe this similarity). Fig. 5B top and bottom panels look visually similar, indicating that the ΔHFD is negatively related to ΔPower.

**Fig. 5:**
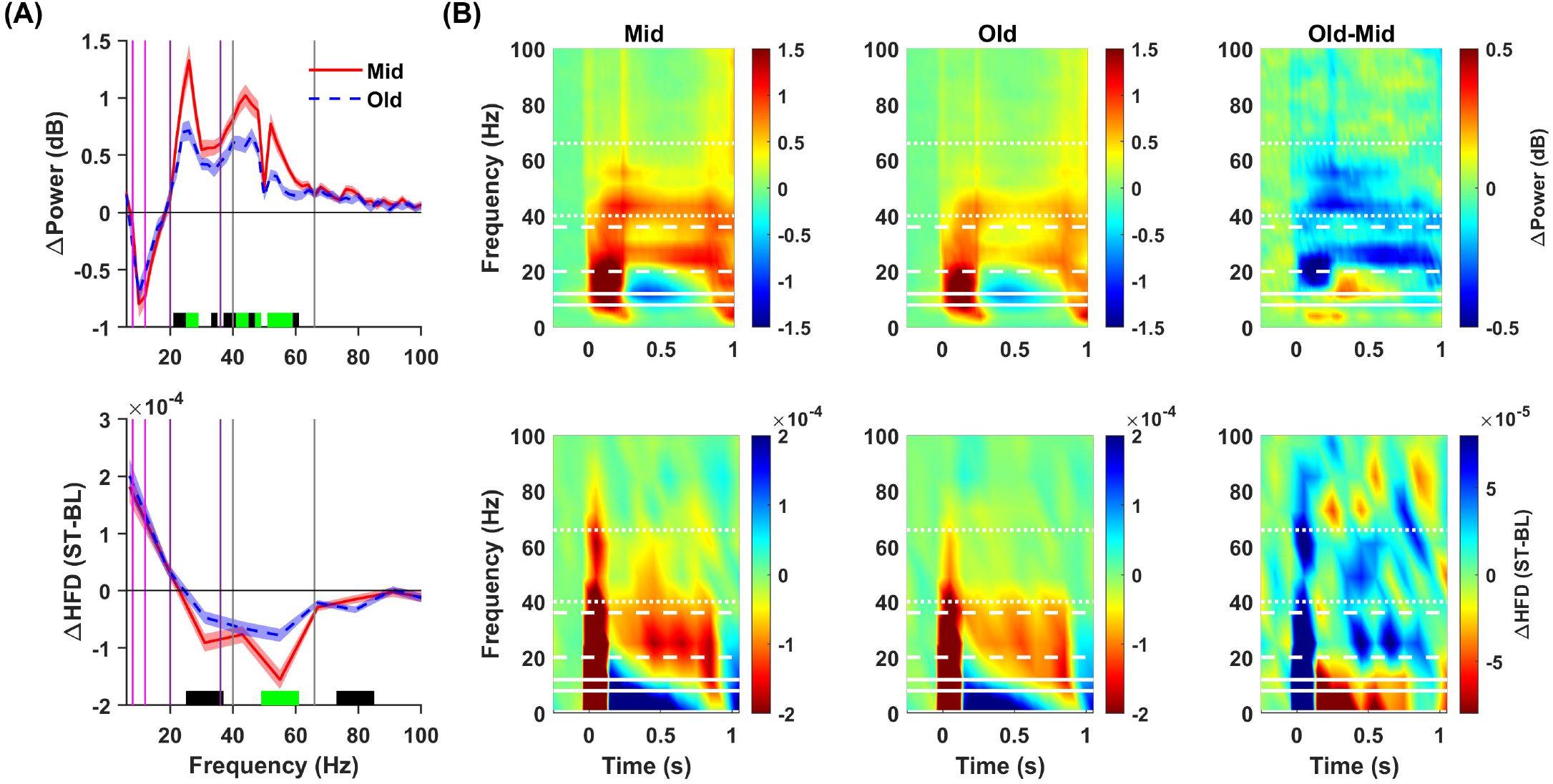
**(A)** Change in power in dB (top) and HFD (ST-BL; bottom) during the ST with respect to the BL for the two age groups. Solid and dash traces represent the median for Mid and Old groups respectively and shaded region around them indicates ±SEM across subjects, computed after bootstrapping over 10,000 iterations. Coloured bars at the abscissa in the bottom panel represent significance of differences of slopes between mid and old (black: p < 0.05 and green: p < 0.01, K-W test). The pink, purple and grey vertical lines represent alpha (8-12 Hz), slow-gamma (20-34 Hz) and fast-gamma (36-66 Hz) frequency bands respectively. **(B)** Delta TF plots for change in power (top) and HFD (bottom) for mid (left), old (centre) and difference (old-mid; right).

#### 3.2.3 Dependence of HFD on age vs spectral measures

To quantify the dependence of HFD on age beyond spectral measures, we sub-selected the subjects from each group such that they had same distribution of slopes and ΔPower in specific bands ((Kumar and Ray, 2023); See Materials and Methods). When we matched subjects for slopes in LFR (64-140 Hz; Fig. 6A left) and HFR (210-290 Hz; Fig. 6A right) respectively, the corresponding difference of HFD in those frequency ranges became non-significant, while the difference in other frequency band remained similar to the non-matched criterion. Similar result held true for the matching of ΔPower in slow gamma (20-34 Hz; Fig. 6B left) and fast gamma (36-66 Hz; Fig. 6B right) frequency ranges. This indicated a strong dependence of age-related HFD changes on the changes in slopes and oscillatory power. We further verified these results using multiple regression analysis that depicted a strong dependence of HFD on spectral measures but not on age.

**Fig. 6:**
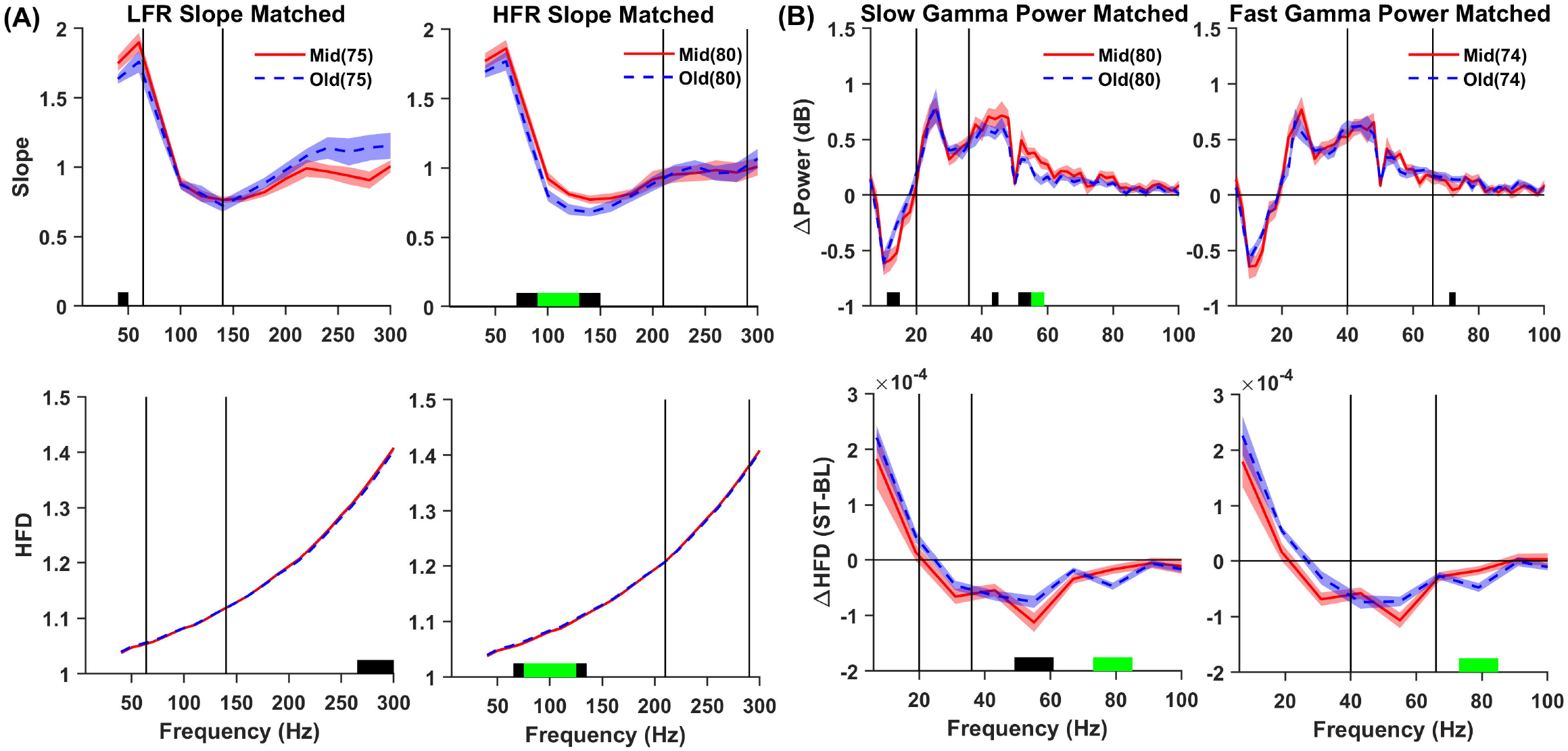
**(A)** BL Slope (top) and HFD (bottom) for slope matched subjects in LFR: 64-140 Hz (Left) and HFR: 210-290 Hz (Right). **(B)** Change in power (top) and the change in HFD (bottom) with respect to the stimulus for Δ Power matched subjects in slow gamma (20-34Hz; left) and fast gamma (36-66 Hz; right) frequency bands. The black vertical lines indicate the corresponding frequency bands where matching is performed. The numbers in the legend indicate the subjects obtained after matching. Solid and dash traces represent the median for Mid and Old groups respectively, and shaded region around them indicates ± SEM across subjects, computed after bootstrapping over 10,000 iterations. Coloured bars at the abscissa in the bottom panel represent significance of differences between mid and old (black: p < 0.05 and green: p < 0.01, K-W test).

**Fig. 7:**
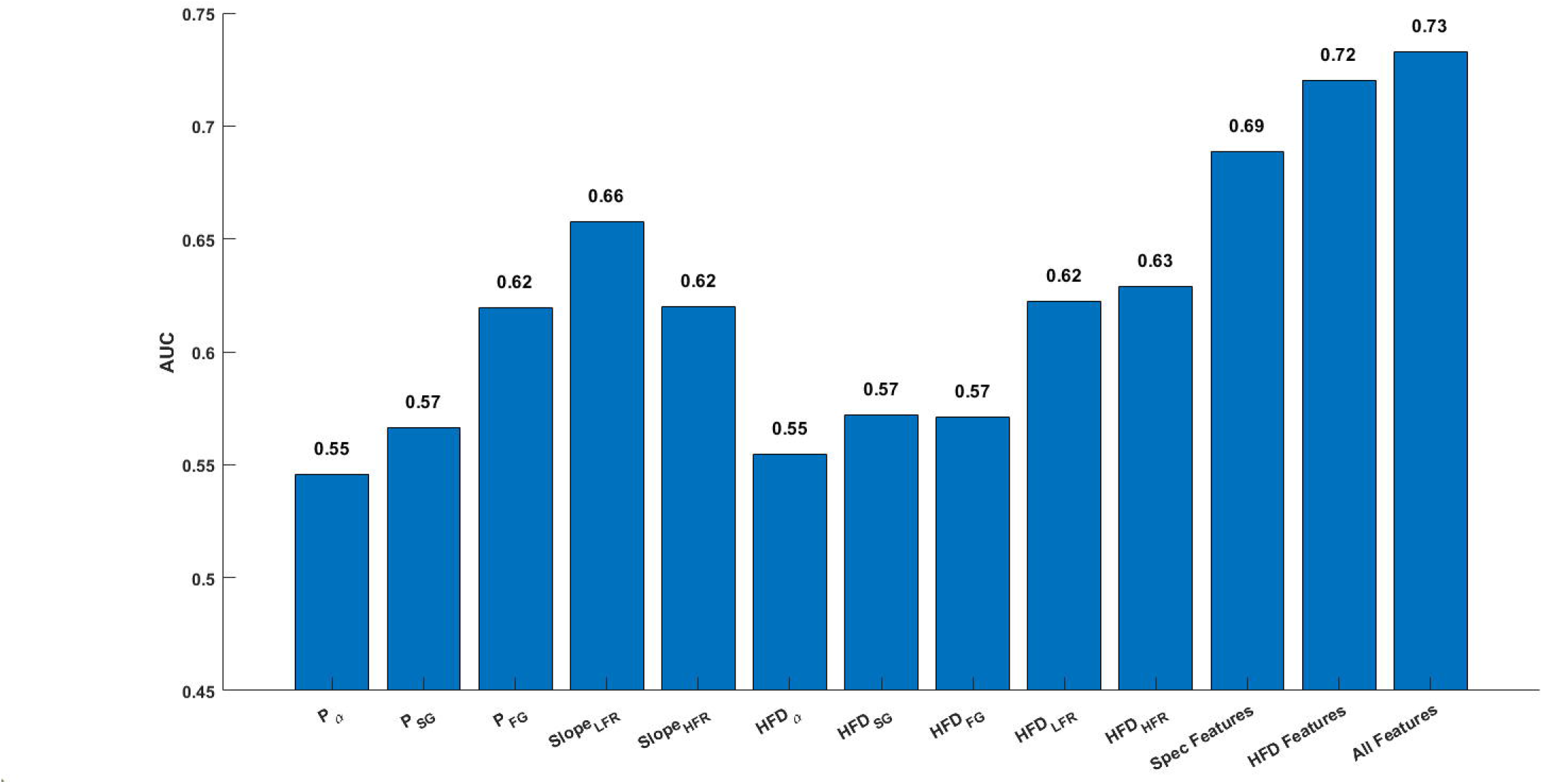
AUC for different spectral and HFD features individually, all spectral features combined as ‘Spec features’, all HFD features combined as ‘HFD features’ and combining all features, computed using 5-fold validation.

### 3.3 Does HFD provide some additional information than spectral features?

#### 3.3.1 Classification using HFD

To test whether HFD provides any extra information about the age than those obtained from spectral features, we explored the classifier performance for age distinctiveness using LDA. We used 5 spectral features: alpha power (*P*_*α*_; 8-12 Hz), slow gamma power (*P*_*SG*_; 20-34 Hz), fast gamma power (*P*_*FG*_; 36 to 66 Hz), slope in low frequency range (*Slope*_*LFR*_; 64-140 Hz) and slope in high frequency range (*Slope*_*LFR*_; 210 to 290 Hz) and 5 HFD features in the respective frequency ranges: *HFD*_*α*_,*HFD*_*SG*_, *HFD*_*FG*_, *HFD*_*LFR*_ and *HFD*_*HFR*_ At individual feature level, *Slope*_*LFR*_ performed the best (AUC = 0.66) among all features. However, after combining spectral and HFD features separately, we found AUC to be higher for all HFD features combined (AUC = 0.72) than all spectral features combined (AUC = 0.69). Adding spectral and HFD features marginally improved the AUC to 0.73. These results generally remained consistent even when this analysis was performed with different choices for frequency ranges for power or slope analysis. Overall, these observations suggest that although HFD and spectral measures were largely redundant, HFD could provide some additional information compared to spectral measures.

#### 3.3.2 Comparison of HFD on real versus surrogate data

To determine whether the changes in HFD with aging or stimulus were just a reflection of linear properties like PSD, we computed the HFD for corresponding surrogate data, which were generated using two different methods: phase randomization and iterated amplitude adjusted Fourier transform (IAAFT; See Materials and Methods). Figure 8A shows the surrogates generated by these two different methods for a sample time series for BL period shown in Fig. 1B. While phase randomised surrogates preserved only PSD of the time series, IAAFT preserved both amplitude distribution and PSD as depicted in Fig. 8B and C. If the HFD only depended on the PSD based properties, we expected the HFDs to be comparable for the three signals. However, the HFDs of phase randomized and IAFFT surrogates were 1.215 and 1.212, which were considerably larger than the HFD of the real signal (1.137). Following (Prichard and Theiler, 1994), we first generated 39 surrogates (for significance level α<0.05; Number of surrogates 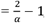 two-sided test, Schreiber and Schmitz, 2000) for the sample time series and computed sigma (S) as 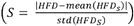 where *HFD*_*S*_ are the HFD for surrogates. The time series is considered non-linear with α<0.05 if S is greater than 2. S for the sample time series was 12.65 for phase randomized and 7.43 for IAAFT surrogates, indicating the presence of non-linearity in EEG. Similar results were observed for the population data.

**Fig. 8:**
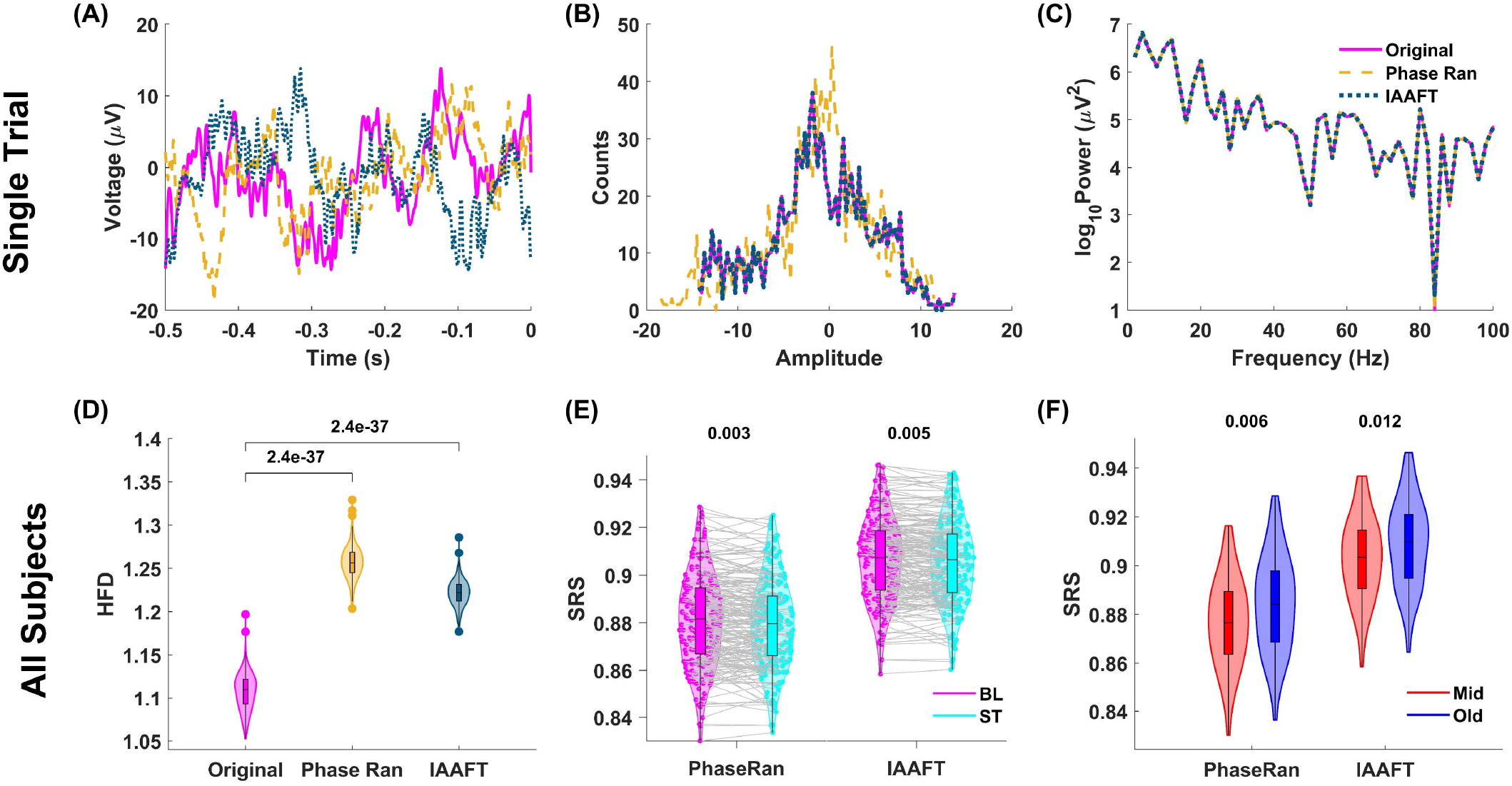
**(A)** Surrogates generated by phase randomization (Phase Ran; dashed) and IAAFT (dotted) for the sample time series for BL period for electrode Oz (Original; solid). The corresponding amplitude distribution and PSD for original and surrogate time series are shown in **(B)** and **(C). (D)** Comparison of median BL HFD for all subjects for signals filtered between 1 and 90 Hz for HP electrode group between original and surrogates. **(E)** and **(F)** shows the comparison of Surrogate Ratio Score (SRS) between ‘BL and ST’ and ‘Mid and Old’ for the surrogates obtained by these two methods respectively. The p-values for median computed from Wilcoxon signed rank sum test for (D) and (E) and WRS test for (F) are indicated on top in each panel.

Next, we generated one surrogate (for time series filtered between 1 and 90 Hz) for each trial and each electrode for all subjects. In HP electrode group for BL, the surrogates had significantly higher HFD as compared to the original data series (Fig. 8D; p=2.4×10^−37^, zval = −12.77, signed rank = 0, Wilcoxon signed rank test for both phase randomization and IAAFT), indicating that there is some non-linear characteristics in EEG that is captured by HFD. Next, we calculated ‘Surrogate Ratio Score’ (SRS), the ratio of original HFD (with both linear and non-linear structures) to the surrogate HFD (with only linear structure), which essentially “normalizes” the PSD based (linear) contribution to HFD (Schartner et al., 2017; Kosciessa et al., 2020). If age or stimulus related changes in HFD were solely due to changes in PSDs, we expect no difference in SRS scores with stimulus or aging. However, SRS was different between BL and ST (Fig. 8E; phase randomization: p = 0.003, zval = 2.94, signed rank = 14551; IAAFT: p = 0.005, zval = 2.79, signed rank = 14411, Wilcoxon signed rank test) as well as mid and old age groups (Fig. 8F; phase randomization: p = 0.006, zval = −2.72, ranksum = 8569; IAAFT: p = 0.012, zval = −2.51, ranksum = 8666, WRS test). It, thereby, shows that changes in HFD with stimulus and age are not solely due to changes in linear properties, and therefore both stimulus and age affect some non-linear dynamics in brain.

## 4. Discussion

We investigated the reflection of brain’s dynamics in HFD and its dependence on the changes in PSD components (oscillations and the slope of aperiodic activity) by studying the age-related variation of HFD in a large (N=217) EEG dataset. The subjects performed a fixation task, where they were shown visual grating stimulus. HFD decreased during the presentation of the stimulus and the reduction was more in mid-aged group (50-65 years) than the old-aged group (>65 years). For BL period, equivalent to eyes open state, HFD increased with age. However, this age-related increase in HFD was limited to frequencies less than 150 Hz and showed opposite trend at higher frequencies. Further, the age-related changes in HFD for the signals, filtered in different frequency range, was negatively related to baseline slope and stimulus-induced oscillatory power in different frequency bands. However, stimulus and age-related changes in HFD persisted even after normalization using surrogate data that preserves the linear properties (power spectrum and amplitude distribution) of the signal, suggesting that changes in non-linear properties with age/stimulus is captured using HFD.

### 4.1 Neural Dynamics reflected in HFD

We found that presentation of a visual grating stimulus lowered the HFD, supporting that HFD may be inversely related to brain synchronicity, similar to what was previously shown for correlation dimension (Stam, 2005). A previous modelling study has shown similar reduction in entropy and complexity due to the stimulus (Ponce-Alvarez et al., 2015). Also, Luczak and colleagues (2009) reported a stimulus-based reduction of multidimensional space outlined by cortical activity. This stimulus-induced decrease in neural variability in terms of reduced firing rate variability has been explained via a spontaneous multi-state stability model (Deco and Hugues, 2012; Litwin-Kumar and Doiron, 2012) and has been linked to information processing. Under the model, the spontaneous activity of local neural networks transits through multiple spontaneous states, thus, rendering it highly heterogeneous (Deco and Hugues, 2012) and has high information capacity (McIntosh et al., 2010). On the other hand, the stimulus stabilizes the network in a single evoked state, resulting in the suppression of the transitions, decrease in variability and enhancement of the transmission of the information about the stimulus (Ponce-Alvarez et al., 2015). However, stimulus onset also suppresses low-frequency oscillations such as alpha or beta, which can increase HFD. For example, increase in HFD has been reported with respect to sound stimuli (Gladun, 2021). Therefore, there can be an increase or decrease in complexity and HFD with respect to stimulus depending on the frequency range for computation of HFD and the emergence or suppression of oscillations. Here, the decrease in HFD during the grating stimulus is due to the emergence of stimulus-induced narrowband gamma oscillations.

The quantification of brain health using HFD is debated in literature. On one hand, the decrease in HFD and thereby, a decrease in complexity due to certain neurodegenerative disorders like Alzheimer’s disease may indicate the decrease in efficiency of neural networks (Smits et al., 2016; Bunimovich and Skums, 2024), thereby, favoring an increase of HFD for a healthy brain. On the other hand, an increase in HFD and hence complexity, like in stress (Phutela et al., 2024) has been related to the reduction in brain synchronicity (Stam, 2005) supporting against the increase of HFD for healthy brain. In the present study, the increase in BL HFD with age could indicate a reduction in long-range neural correlations with aging as HFD is negatively associated with H, a measure of long range correlations (Hurst, 1951; Shang, 2020).

The results in the present work seems inconsistent with the previous works (Zappasodi et al., 2015; Smits et al., 2016), who showed an inverse parabola relation of HFD with age between 20-80 years with the decrease in HFD above 50 years. However, HFD values are critically dependent on the choice of *k*_*max*_. While Zappasodi and colleagues (2015) chose *k*_*max*_ 16, Smits and colleagues (2016) had chosen it as 65 and reported a decrease in HFD with age for elderly population. However, the latter work found an increase in HFD for older population at low *k*_*max*_ (Fig 2. in (Smits et al., 2016)). Another study also reported an increase in correlation dimension with age, albeit over a different age range of 20-60 years (Anokhin et al., 1996).

### 4.2 Frequency dependence of HFD

The increase in HFD with frequency has also been observed previously but up to 47 Hz only (Ferenets et al., 2007). This is mainly due to increase in roughness or irregularity in signal with increase in its frequency. Further, it also supports that long-range correlations are difficult to maintain at high frequencies. Differences in age-related variation in HFD in low (<150 Hz) and high frequency ranges (> 150 Hz) indicates the role of frequency range in governing the changes in HFD. We did EEG analysis beyond the commonly chosen range as in our previous work, we found an opposite age-related slope change in high frequency range as compared to the traditional frequency range (Aggarwal and Ray, 2023). High frequency (>150 Hz) analysis of EEG may provide information about the neural processes with time scales less than 10 ms like synaptic neurotransmitter diffusion and spike propagation (Sabatini and Regehr, 1996; Shepherd, 2004).

### 4.3 Correlation of HFD with spectral measures

The change in HFD was strongly negatively correlated with change in slope (Fig. 4C right bottom panel). While slope reflects linear spectral properties of the signal, HFD is a measure of non-linear complexity. Conceptually, these metrics are designed to capture different aspects of neural dynamics, so such a robust correlation (−0.99) between these measures was surprising. It may indicate towards a systematic relationship between linear and non-linear properties in brain signals, possibly reflecting a shared underlying mechanism or coupling between the spectral and temporal complexities of neural activity. However, the exact nature of this relationship remains unclear, and warrants further investigation in future studies.

Despite of the presence of the negative correlation between ‘changes’ in slope and HFD, we did not find a straightforward dependence of HFD on slope (Fig. 4C left bottom panel), unlike demonstrated for synthetic signals (Higuchi, 1990) and for colored noise in Fig. 1A. This may be due to our choice of fixing *k*_*max*_ to the same value of 5 for all frequency ranges. Also, 1/f slope could be modified due to the presence of oscillations and the intrinsic variation in PSD (Aggarwal and Ray, 2023; Merkin et al., 2023), which could influence the fall of the PSD. Therefore, the negative correlation became visible only with respect to changes in HFD and slope after the inherent noise cancellation.

Interestingly, changes in HFD were strongly dependent on the changes in slope rather than the change in broadband power (Fig. 4C right panels), even though, both higher broadband power and shallower slopes at high frequencies are indicators of greater noise in a system. This could be due to different neural mechanisms controlling the origin of broadband power and slopes. For example, while broadband power in high-gamma range is thought to be generated due to increased spiking activity in microelectrode recordings (Ray and Maunsell, 2011), E-I balance is among the widely accepted reasons for the slope modulation (Gao et al., 2017). Though speculative, the dependence of HFD on slope may indicate that HFD could better reflect E-I balance. Moreover, E-I balance plays an important role in the generation of stimulus-induced oscillatory gamma power (Wilson and Cowan, 1972; Fuchs et al., 2007; Buzsáki and Wang, 2012). Thus, the negative dependence of HFD on stimulus-induced gamma power (Fig. 6) provides further hints for the existence of a relation between HFD and E-I balance. Future studies can help to shed light on more reasons for the dependence of HFD on slope and stimulus-induced gamma power but not overall broadband power.

We found conflicting description about the correlation between HFD and power in different frequency bands in literature. This could be because power in a given frequency band may increase due to an oscillation in that frequency range (bump in the PSD) or an increase in broadband power which leads to an increase in the “pedestal” of the PSD, even though these two types of change (narrowband versus broadband) may have opposite effects on HFD. For example, HFD has been shown to decrease in chronic insomniacs compared to controls, which was related to increased alpha power, mainly in the right hemisphere, in patients with insomnia (Kronholm et al., 2007). Further, HFD has been correlated negatively with theta activity but positively with gamma activity (Stojadinović et al., 2020). Zappasodi and colleagues (2015) had shown negative correlations of HFD with power in theta and alpha ranges. Along with direct comparisons, indirect correlations may be inferred by comparing HFD and power-based studies. For example, in depression, both HFD (Akar et al., 2015) and power (Strelets et al., 2007) in gamma band increases, indicating a positive correlation. Similarly, reduced HFD in AD (Smits et al., 2016) and increased gamma power during resting state (van Deursen et al., 2008; Güntekin et al., 2020; Fide et al., 2021) hint at a negative correlation. P □ eske and colleagues surveyed several studies and found HFD to be negatively correlated with theta, alpha and beta band power but weakly positively related to gamma band power (Päeske et al., 2023). However, most of these studies involved resting state brain activity (Kronholm et al., 2007; Smits et al., 2016; Stojadinović et al., 2020), that is dominated by low frequency peak oscillations, so their high frequency activity may not be oscillatory. Also, in these studies, HFD computed for broadband frequency ranges were correlated with narrow band oscillations, while we computed HFD and power in the same frequency bands. An important point is that during the BL period in the present study (Fig. 4), the old-aged group has flatter PSD and corresponding higher power above 50 Hz. If we consider power in gamma range (30-80 Hz), our results would also indicate that increase in HFD is related to increase in broadband gamma power, which is different from stimulus-induced narrowband gamma oscillations described earlier. So, one needs to be cautious about the oscillatory power that are characterised by peaks in the PSD spectrum and the broadband power that can be indicative of slopes.

### 4.4 HFD as a classifier

Classification studies provide mixed results about the performance comparison between HFD and band power. While HFD and band power performed equally for the classification of schizophrenic patients (Sabeti et al., 2007) and depressive state using LDA classifier (Hosseinifard et al., 2013), HFD showed slightly higher accuracy (3.3%) over alpha band power for the latter classification when a different classification technique (logistic regression) was used (Hosseinifard et al., 2013). These results indicate a strong dependence of HFD on power similar to the present work where HFD performed slightly better than power measures. However, the improvement of AUC using HFD by ~0.03 (3%; AUC_HFD_ = 0.72 and AUC_Spec_ = 0.69) hints at an advantage of nonlinear methods over linear methods for classification.

### 4.5 Reflection of non-linear brain dynamics in HFD via surrogate analysis

Difference in HFD between the original EEG time series and their surrogates indicates the presence of non-linearity in brain signals. Though HFD is dependent on linear PSD changes, it also captures the changes in non-linear dynamics in brain due to aging as well as stimulus. There are very few age-related studies that involve surrogate analysis while using non-linear techniques. While (Müller and Lindenberger, 2012) found differences between real and surrogate data using pointwise dimension analysis for all age groups, other studies (Courtiol et al., 2016; Kosciessa et al., 2020) have reported that age-related differences in multi-scale entropy were mainly due to linear correlations. Though non-linear dynamics has been shown to alter across sleep states (Schartner et al., 2017), the non-linear effect of aging using surrogate analysis has not been reported previously to our knowledge.

In conclusion, while there is a clear dependence of age-related HFD changes on spectral measures (including oscillatory power and slope) in neural signals, this does not mean they are merely different representations of the same underlying information. Classification results show that HFD contributes unique information related to age. Additionally, surrogate analysis demonstrates that HFD captures some non-linear dynamics of neural signals. This observed dependence may reflect an underlying relationship between linear and non-linear aspects of brain activity. Overall, our findings highlight the need for new models that can jointly account for both linear and non-linear characteristics of brain signals.

## Abbreviations

EEG: Electroencephalogram
PSD: Power Spectral Density
FD: Fractal Dimension
HFD: Higuchi’s Fractal Dimension
H: Hurst Index
CD: Correlation Dimension
E-I: Excitation-Inhibition
LDA: Linear Discriminant Analysis
fBm: fractional Brownian motion
NIMHANS: National Institute of Mental Health and Neurosciences
ACE-III: Addenbrooke’s Cognitive Examination-III
CDR: Clinical Dementia Rating
HMSE: Hindi Mental State Examination
SD: Standard Deviation
SEM: Standard Error of Median
RMS: Root Mean Square
BL: Baseline
ST: Stimulus
FOOOF: Fitting Oscillations and One Over f
ROC: Receiver Operating Characteristic
AUC: Area under the ROC curve
IAAFT: Iterated Amplitude Adjusted Fourier Transform
HP: High-Priority
K-W: Kruskal-Wallis
WRS: Wilcoxon rank sum
FDR: False Discovery Rate
O: Occipital
CP: Centro-Parietal
FC: Fronto-Central
F: Frontal
T: Temporal
S: Sigma
SRS: Surrogate Ratio Score

## References

Accardo A, Affinito M, Carrozzi M, Bouquet F (1997) Use of the fractal dimension for the analysis of electroencephalographic time series. Biol Cybern 77:339–350.

Aggarwal S, Ray S (2023) Slope of the power spectral density flattens at low frequencies (<150 Hz) with healthy aging but also steepens at higher frequency (>200 Hz) in human electroencephalogram. Cerebral Cortex Communications 4:tgad011.

Akar SA, Kara S, Agambayev S, Bilgic V (2015) Nonlinear analysis of EEG in major depression with fractal dimensions. Annu Int Conf IEEE Eng Med Biol Soc 2015:7410–7413.

Amezquita-Sanchez JP, Mammone N, Morabito FC, Marino S, Adeli H (2019) A novel methodology for automated differential diagnosis of mild cognitive impairment and the Alzheimer’s disease using EEG signals. Journal of Neuroscience Methods 322:88–95.

Anokhin AP, Birbaumer N, Lutzenberger W, Nikolaev A, Vogel F (1996) Age increases brain complexity. Electroencephalography and Clinical Neurophysiology 99:63–68.

Babiloni C, Binetti G, Cassarino A, Dal Forno G, Del Percio C, Ferreri F, Ferri R, Frisoni G, Galderisi S, Hirata K, Lanuzza B, Miniussi C, Mucci A, Nobili F, Rodriguez G, Luca Romani G, Rossini PM (2006) Sources of cortical rhythms in adults during physiological aging: a multicentric EEG study. Hum Brain Mapp 27:162–172.

Barabási A-L, Vicsek T (1991) Multifractality of self-affine fractals. Phys Rev A 44:2730–2733.

Benjamini Y, Hochberg Y (1995) Controlling the false discovery rate: a practical and powerful approach to multiple testing. J Roy Statist Soc Ser B 57:289–300.

Bokil H, Andrews P, Kulkarni JE, Mehta S, Mitra PP (2010) Chronux: a platform for analyzing neural signals. J Neurosci Methods 192:146–151.

Bunimovich L, Skums P (2024) Fractal networks: Topology, dimension, and complexity. Chaos: An Interdisciplinary Journal of Nonlinear Science 34:042101.

Buzsáki G, Wang X-J (2012) Mechanisms of Gamma Oscillations. Annu Rev Neurosci 35:203–225.

Churchland MM et al. (2010) Stimulus onset quenches neural variability: a widespread cortical phenomenon. Nat Neurosci 13:369–378.

Courtiol J, Perdikis D, Petkoski S, Müller V, Huys R, Sleimen-Malkoun R, Jirsa VK (2016) The multiscale entropy: Guidelines for use and interpretation in brain signal analysis. Journal of Neuroscience Methods 273:175–190.

Deco G, Hugues E (2012) Neural Network Mechanisms Underlying Stimulus Driven Variability Reduction. PLOS Computational Biology 8:e1002395.

Delorme A, Makeig S (2004) EEGLAB: an open source toolbox for analysis of single-trial EEG dynamics including independent component analysis. J Neurosci Methods 134:9–21.

Di Ieva A ed. (2024) The Fractal Geometry of the Brain. Cham: Springer International Publishing. Available at: https://link.springer.com/10.1007/978-3-031-47606-8 [Accessed November 19, 2024].

Díaz Beltrán L, Madan CR, Finke C, Krohn S, Di Ieva A, Esteban FJ (2024) Fractal Dimension Analysis in Neurological Disorders: An Overview. In: The Fractal Geometry of the Brain (Di Ieva A, ed), pp 313–328. Cham: Springer International Publishing. Available at: 10.1007/978-3-031-47606-8_16 [Accessed May 6, 2024].

Donoghue T, Haller M, Peterson EJ, Varma P, Sebastian P, Gao R, Noto T, Lara AH, Wallis JD, Knight RT, Shestyuk A, Voytek B (2020) Parameterizing neural power spectra into periodic and aperiodic components. Nat Neurosci 23:1655–1665.

Drachman DA (2006) Aging of the brain, entropy, and Alzheimer disease. Neurology 67:1340–1352.

Eke A, Hermán P, Bassingthwaighte J, Raymond G, Percival D, Cannon M, Balla I, Ikrényi C (2000) Physiological time series: distinguishing fractal noises from motions. Pflügers Arch – Eur J Physiol 439:403–415.

Esteller R, Vachtsevanos G, Echauz J, Litt B (2001) A comparison of waveform fractal dimension algorithms. IEEE Trans Circuits Syst I 48:177–183.

Falconer K (2013) Fractal Geometry: Mathematical Foundations and Applications. John Wiley & Sons.

Ferenets R, Vanluchene A, Lipping T, Heyse B, Struys MMRF (2007) Behavior of entropy/complexity measures of the electroencephalogram during propofol-induced sedation: dose-dependent effects of remifentanil. Anesthesiology 106:696–706.

Fide E, Yerlikaya D, Yener G (2021) Resting-state EEG gamma power and coherence might be an indicator of hyperexitability in patients with early-onset Alzheimer’s disease. Alzheimer’s & Dementia 17:e058669.

Fuchs EC, Zivkovic AR, Cunningham MO, Middleton S, LeBeau FEN, Bannerman DM, Rozov A, Whittington MA, Traub RD, Rawlins JNP, Monyer H (2007) Recruitment of Parvalbumin-Positive Interneurons Determines Hippocampal Function and Associated Behavior. Neuron 53:591–604.

Gao R, Peterson EJ, Voytek B (2017) Inferring synaptic excitation/inhibition balance from field potentials. NeuroImage 158:70–78.

Gladun KV (2021) Higuchi Fractal Dimension as a Method for Assessing Response to Sound Stimuli in Patients with Diffuse Axonal Brain Injury. Sovrem Tekhnologii Med 12:63–70.

Güntekin B, Yıldırım E, Kıyı İ, Fide E, Uzunlar H, Calısoglu P, Yirikogullari H, Akturk T, Yener G (2020) Gamma power increased in Alzheimer’s disease patients in comparison to healthy controls during recognition of facial expressions. Alzheimer’s & Dementia 16:e043387.

Harada CN, Natelson Love MC, Triebel KL (2013) Normal Cognitive Aging. Clinics in Geriatric Medicine 29:737–752.

Higuchi T (1988) Approach to an irregular time series on the basis of the fractal theory. Physica D: Nonlinear Phenomena 31:277–283.

Higuchi T (1990) Relationship between the fractal dimension and the power law index for a time series: A numerical investigation. Physica D: Nonlinear Phenomena 46:254–264.

Hogan MJ, Kilmartin L, Keane M, Collins P, Staff RT, Kaiser J, Lai R, Upton N (2012) Electrophysiological entropy in younger adults, older controls and older cognitively declined adults. Brain Research 1445:1–10.

Hosseinifard B, Moradi MH, Rostami R (2013) Classifying depression patients and normal subjects using machine learning techniques and nonlinear features from EEG signal. Computer Methods and Programs in Biomedicine 109:339–345.

Hurst HE (1951) Long-Term Storage Capacity of Reservoirs. Transactions of the American Society of Civil Engineers 116:770–799.

Kakumanu RJ, Nair AK, Venugopal R, Sasidharan A, Ghosh PK, John JP, Mehrotra S, Panth R, Kutty BM (2018) Dissociating meditation proficiency and experience dependent EEG changes during traditional Vipassana meditation practice. Biological Psychology 135:65–75.

Kesić S, Spasić SZ (2016) Application of Higuchi’s fractal dimension from basic to clinical neurophysiology: A review. Computer Methods and Programs in Biomedicine 133:55–70.

Kosciessa JQ, Kloosterman NA, Garrett DD (2020) Standard multiscale entropy reflects neural dynamics at mismatched temporal scales: What’s signal irregularity got to do with it? PLOS Computational Biology 16:e1007885.

Kronholm E, Virkkala J, Kärki T, Karjalainen P, Lang H, Hämäläinen H (2007) Spectral power and fractal dimension: Methodological comparison in a sample of normal sleepers and chronic insomniacs. Sleep Biol Rhythms 5:239–250.

Kumar WS, Manikandan K, Murty DVPS, Ramesh RG, Purokayastha S, Javali M, Rao NP, Ray S (2022) Stimulus-Induced Narrowband Gamma Oscillations are Test–Retest Reliable in Human EEG. Cereb Cortex Commun 3:tgab066.

Kumar WS, Ray S (2023) Healthy ageing and cognitive impairment alter EEG functional connectivity in distinct frequency bands. European Journal of Neuroscience 58:3432–3449.

Lau ZJ, Pham T, Chen SHA, Makowski D (2022) Brain entropy, fractal dimensions and predictability: A review of complexity measures for EEG in healthy and neuropsychiatric populations. European Journal of Neuroscience 56:5047–5069.

Litwin-Kumar A, Doiron B (2012) Slow dynamics and high variability in balanced cortical networks with clustered connections. Nat Neurosci 15:1498–1505.

Luczak A, Barthó P, Harris KD (2009) Spontaneous Events Outline the Realm of Possible Sensory Responses in Neocortical Populations. Neuron 62:413–425.

Mandelbrot BB (1985) Self-Affine Fractals and Fractal Dimension. Phys Scr 32:257.

Mandelbrot BB (2013) Fractals and Scaling in Finance: Discontinuity, Concentration, Risk. Selecta Volume E. Springer Science & Business Media.

Mandelbrot BB, Van Ness JW (1968) Fractional Brownian Motions, Fractional Noises and Applications. SIAM Rev 10:422–437.

Matteo TD, Aste T, Dacorogna MM (2005) Long-term memories of developed and emerging markets: Using the scaling analysis to characterize their stage of development. Journal of Banking & Finance 29:827–851.

McIntosh AR, Vakorin V, Kovacevic N, Wang H, Diaconescu A, Protzner AB (2014) Spatiotemporal Dependency of Age-Related Changes in Brain Signal Variability. Cerebral Cortex 24:1806–1817.

Merkin A, Sghirripa S, Graetz L, Smith AE, Hordacre B, Harris R, Pitcher J, Semmler J, Rogasch NC, Goldsworthy M (2023) Do age-related differences in aperiodic neural activity explain differences in resting EEG alpha? Neurobiology of Aging 121:78–87.

Molz FJ, Liu HH, Szulga J (1997) Fractional Brownian motion and fractional Gaussian noise in subsurface hydrology: A review, presentation of fundamental properties, and extensions. Water Resources Research 33:2273–2286.

Morel P (2018) Gramm: grammar of graphics plotting in Matlab. Journal of Open Source Software 3:568.

Müller V, Lindenberger U (2012) Lifespan differences in nonlinear dynamics during rest and auditory oddball performance. Developmental Science 15:540–556.

Murty DV, Manikandan K, Kumar WS, Ramesh RG, Purokayastha S, Nagendra B, Ml A, Balakrishnan A, Javali M, Rao NP, Ray S (2021) Stimulus-induced gamma rhythms are weaker in human elderly with mild cognitive impairment and Alzheimer’s disease Vinck M, Colgin LL, Bosman CA, eds. eLife 10:e61666.

Murty DVPS, Manikandan K, Kumar WS, Ramesh RG, Purokayastha S, Javali M, Rao NP, Ray S (2020) Gamma oscillations weaken with age in healthy elderly in human EEG. NeuroImage 215:116826.

Murty DVPS, Ray S (2022) Stimulus-induced Robust Narrow-band Gamma Oscillations in Human EEG Using Cartesian Gratings. Bio Protoc 12:e4379.

Olejarczyk E, Cukic M, Porcaro C, Zappasodi F, Tecchio F (2024) Clinical Sensitivity of Fractal Neurodynamics. In: The Fractal Geometry of the Brain (Di Ieva A, ed), pp 285–312. Cham: Springer International Publishing.

Orey S (1970) Gaussian sample functions and the Hausdorff dimension of level crossings. Z Wahrscheinlichkeitstheorie verw Gebiete 15:249–256.

Päeske L, Uudeberg T, Hinrikus H, Lass J, Bachmann M (2023) Correlation between electroencephalographic markers in the healthy brain. Sci Rep 13:6307.

Phillips LH, Andrés P (2010) The cognitive neuroscience of aging: new findings on compensation and connectivity. Cortex; a Journal Devoted to the Study of the Nervous System and Behavior 46:421–424.

Phutela N, Gabrani G, Kumaraguru P, Relan D (2024) Effectiveness of Higuchi fractal dimension in differentiating subgroups of stressed and non-stressed individuals. Multimed Tools Appl 83:52433–52450.

Ponce-Alvarez A, He BJ, Hagmann P, Deco G (2015) Task-Driven Activity Reduces the Cortical Activity Space of the Brain: Experiment and Whole-Brain Modeling. PLOS Computational Biology 11:e1004445.

Prichard D, Theiler J (1994) Generating surrogate data for time series with several simultaneously measured variables. Phys Rev Lett 73:951–954.

Rabinovich MI, Varona P, Selverston AI, Abarbanel HDI (2006) Dynamical principles in neuroscience. Rev Mod Phys 78:1213–1265.

Rapp PE, Albano AM, Zimmerman ID, Jiménez-Montaño MA (1994) Phase-randomized surrogates can produce spurious identifications of non-random structure. Physics Letters A 192:27–33.

Ray S, Maunsell JHR (2011) Different Origins of Gamma Rhythm and High-Gamma Activity in Macaque Visual Cortex. PLOS Biology 9:e1000610.

Sabatini BL, Regehr WG (1996) Timing of neurotransmission at fast synapses in the mammalian brain. Nature 384:170–172.

Sabeti M, Boostani R, Katebi SD, Price GW (2007) Selection of relevant features for EEG signal classification of schizophrenic patients. Biomedical Signal Processing and Control 2:122–134.

Scally B, Burke MR, Bunce D, Delvenne J-F (2018) Resting-state EEG power and connectivity are associated with alpha peak frequency slowing in healthy aging. Neurobiol Aging 71:149–155.

Schartner MM, Pigorini A, Gibbs SA, Arnulfo G, Sarasso S, Barnett L, Nobili L, Massimini M, Seth AK, Barrett AB (2017) Global and local complexity of intracranial EEG decreases during NREM sleep. Neuroscience of Consciousness 2017:iw022.

Schreiber T, Schmitz A (1996) Improved Surrogate Data for Nonlinearity Tests. Phys Rev Lett 77:635–638.

Schreiber T, Schmitz A (2000) Surrogate time series. Physica D: Nonlinear Phenomena 142:346–382.

Shang HL (2020) A Comparison of Hurst Exponent Estimators in Long-range Dependent Curve Time Series. Journal of Time Series Econometrics 12 Available at: https://www.degruyter.com/document/doi/10.1515/jtse-2019-0009/html [Accessed November 10, 2024].

Shepherd GM (2004) The Synaptic Organization of the Brain. Oxford University Press. Available at: https://global.oup.com/academic/product/the-synaptic-organization-of-the-brain-9780195159561?cc=in&lang=en& [Accessed February 12, 2023].

Smith JH, Rowland C, Harland B, Moslehi S, Montgomery RD, Schobert K, Watterson WJ, Dalrymple-Alford J, Taylor RP (2021) How neurons exploit fractal geometry to optimize their network connectivity. Sci Rep 11:2332.

Smits FM, Porcaro C, Cottone C, Cancelli A, Rossini PM, Tecchio F (2016) Electroencephalographic Fractal Dimension in Healthy Ageing and Alzheimer’s Disease. PLOS ONE 11:e0149587.

Spasić S, Kalauzi A, Ćulić M, Grbić G, Martać L (2005) Estimation of Parameter kmax in Fractal Analysis of Rat Brain Activity. Annals of the New York Academy of Sciences 1048:427–429.

Stam CJ (2005) Nonlinear dynamical analysis of EEG and MEG: Review of an emerging field. Clinical Neurophysiology 116:2266–2301.

Stojadinović G, Martać L, Podgorać J, Spasić SZ, Petković B, Sekulić S, Kesić S (2020) The effects of Nembutal on the intracerebellar EEG activity revealed by spectral and fractal analysis. Archives of Biological Sciences 72:425–432.

Strelets VB, Garakh ZhV, Novototskii-Vlasov VYu (2007) Comparative study of the gamma rhythm in normal conditions, during examination stress, and in patients with first depressive episode. Neurosci Behav Physiol 37:387–394.

Theiler J, Eubank S, Longtin A, Galdrikian B, Doyne Farmer J (1992) Testing for nonlinearity in time series: the method of surrogate data. Physica D: Nonlinear Phenomena 58:77–94.

van Deursen JA, Vuurman EFPM, Verhey FRJ, van Kranen-Mastenbroek VHJM, Riedel WJ (2008) Increased EEG gamma band activity in Alzheimer’s disease and mild cognitive impairment. J Neural Transm (Vienna) 115:1301–1311.

Voytek B, Kramer MA, Case J, Lepage KQ, Tempesta ZR, Knight RT, Gazzaley A (2015) Age-Related Changes in 1/f Neural Electrophysiological Noise. Journal of Neuroscience 35:13257–13265.

Werner G (2010) Fractals in the Nervous System: Conceptual Implications for Theoretical Neuroscience. Front Physiol 1:15.

Wilson HR, Cowan JD (1972) Excitatory and Inhibitory Interactions in Localized Populations of Model Neurons. Biophysical Journal 12:1–24.

Zappasodi F, Marzetti L, Olejarczyk E, Tecchio F, Pizzella V (2015) Age-Related Changes in Electroencephalographic Signal Complexity. PLOS ONE 10:e0141995.

