## Supplementary Data for "Changes in Higuchi’s fractal dimension across age in healthy human EEG are anti-correlated with changes in oscillatory power and 1/f slope"

**Computation of H**

We used generalised Hurst exponent method to compute H, that involves the q^th^ order moments $K_{q}\left( r \right)$ of the distribution of the increments (Barabási and Vicsek, 1991; Matteo et al., 2005; Mandelbrot, 2013).

$K_{q}(r)$ for the time series $X\left( 1 \right), X\left( 2 \right), X\left( 3 \right),\ldots, X(N)$, is given by

$$K_{q}\left( r \right)=\frac{<\left| X\left( t+r \right)-X\left( t \right) \right|^{q}>}{<\left| X\left( t \right) \right|^{q}>}$$

where *r* varies from 1 to $r_{max}$ (*r* plays a similar role for calculating H as *k* in HFD). $K_{q}\left( r \right)$ is assumed to follow the relation

$$K_{q}\left( r \right)\sim r^{qH\left( q \right)}$$

where *H(q)* is the generalised H.

We used the matlab code (<https://in.mathworks.com/matlabcentral/fileexchange/30076-generalized-hurst-exponent>) to calculate H for first order moments with the default parameters, $q=1$ and $r_{\max}$=19.

We found a close match between theoretical result of FD = 2-H and the observed results, where HFD+H =1.985$\pm$0.001, computed for the experimental data.

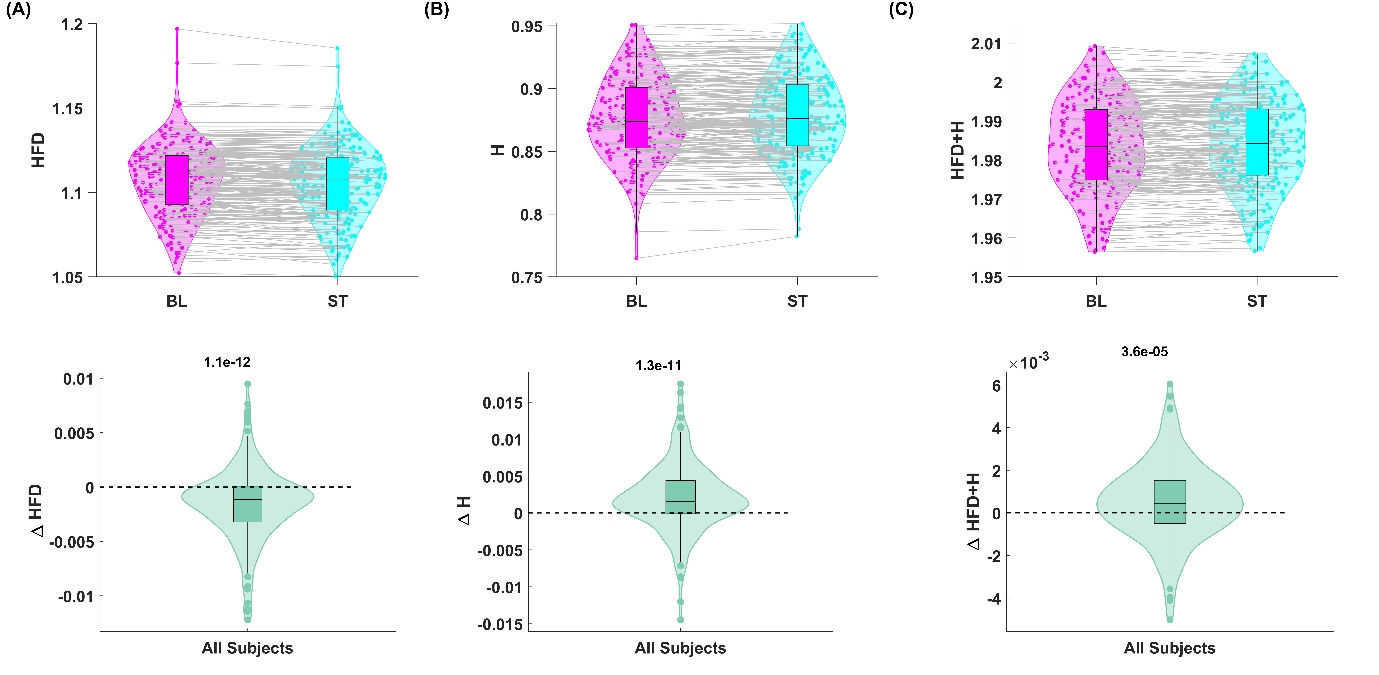

**Supplementary Fig. 1** Violin plots showing the distribution of **(A)** HFD, **(B)** H and **(C)** HFD+H for BL and ST (top panel) and their difference (bottom panel) for all subjects (N=217). p-values obtained from Wilcoxon signed rank test are indicated at the top of the bottom panels.

**Supplementary Tables**

(A) **Baseline**

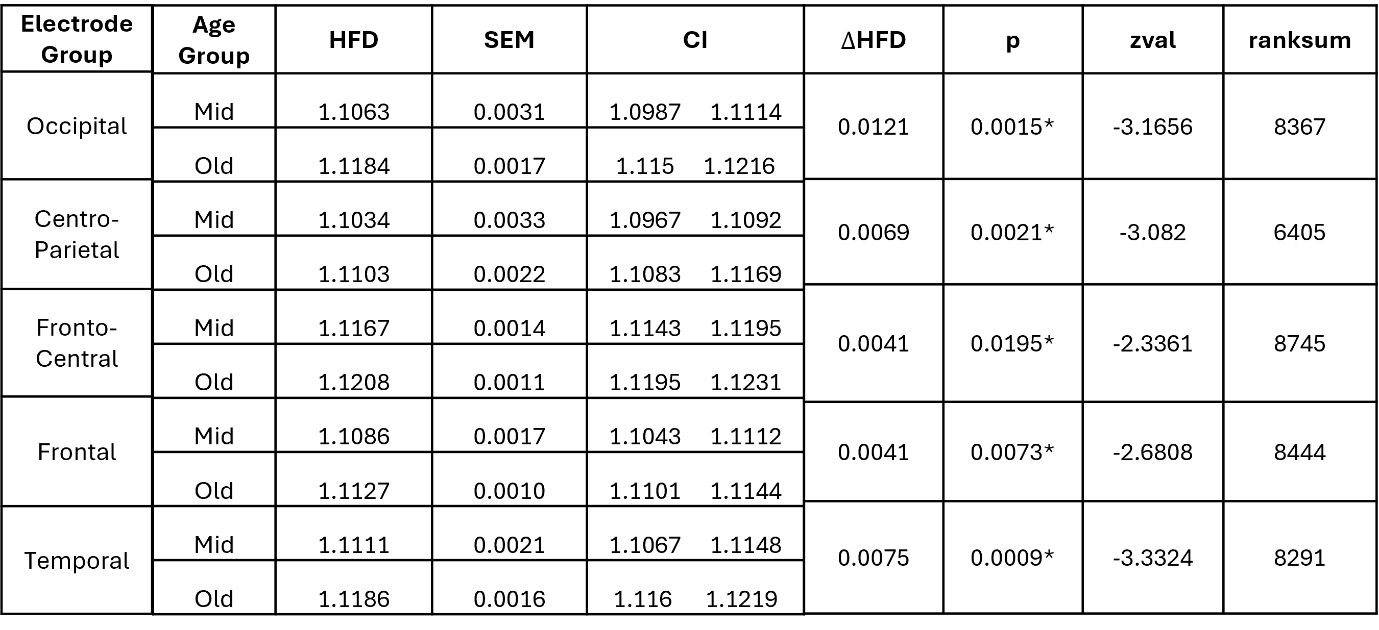

(B) **Stimulus**

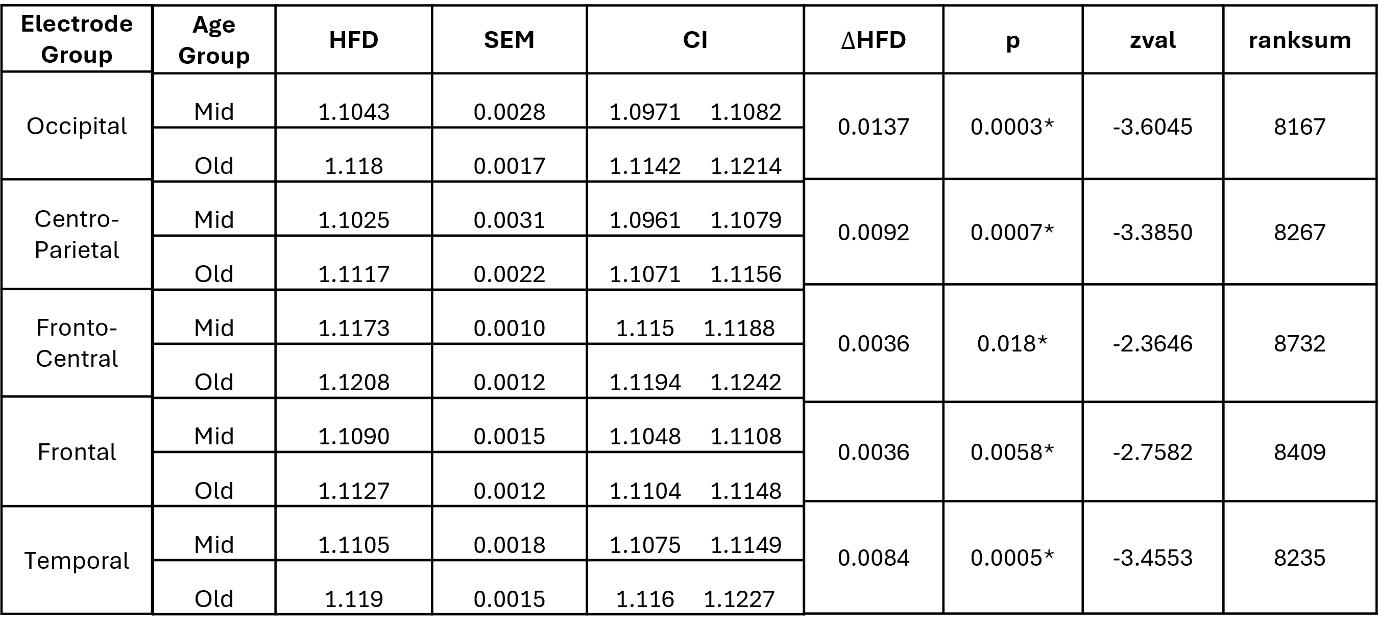

(C) **Stimulus – Baseline (Diff)**

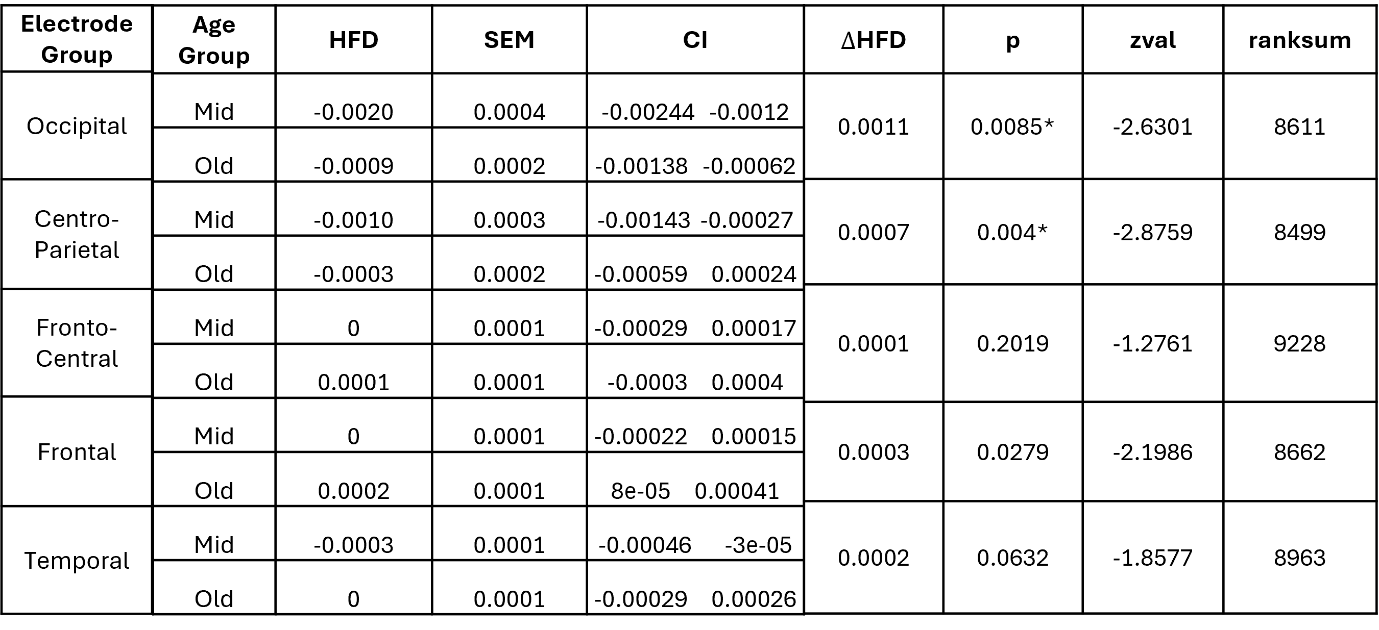

**Supplementary Table 1.** Median HFD, Standard error of median (SEM), 95% Confidence Interval (CI) for the two age groups and their comparison statistics: $\Delta$HFD (difference of the median of Old and Mid, Old-Mid), p-value (* indicates that it is significant after FDR correction), z value (zval) and ranksum for Wilcoxon ranksum test for **(A)** Baseline, **(B)** Stimulus and **(C)** Difference of stimulus and baseline.
